# Immuno-targeting of Gram-positive Pathogens *via* a Cell Wall Binding Tick Antifreeze Protein

**DOI:** 10.1101/2022.09.02.506389

**Authors:** Brianna E. Dalesandro, Marcos M. Pires

## Abstract

The human immune system employs several mechanisms to defend against pathogenic bacteria. However, pathogenic bacterial cells have evolved means to counter these responses, rendering our immune system less effective. Immunological agents that supplement or modulate the host immune response have proven to have powerful therapeutic potential, although this modality is less explored against bacterial pathogens. We describe the application of a bacterial binding protein to re-engage the immune system towards pathogenic bacteria. More specifically, a hapten was conjugated to a protein expressed by *Ixodes scapularis* ticks, called *Ixodes scapularis* antifreeze glycoprotein (IAFGP), that has high affinity for the D-alanine residue on the peptidoglycan of the bacterial cell wall. We showed that a fragment of this protein retained high surface binding affinity. Moreover, when conjugated to a hapten this conjugate led to the display of haptens on the cell surface of vancomycin-resistant *Enterococcus faecalis*. Hapten display then induced the recruitment of antibodies and promoted immune-cell mediated uptake of bacterial pathogens. These results demonstrate the feasibility in using cell wall binding agents as the basis of a class of bacterial immunotherapies against bacterial pathogens.

## Introduction

Findings from over 100 years ago by Emil von Behring demonstrated that a patient’s immune system is, in fact, potent enough to reverse the course of severe bacterial infections (including late stage disease).^1^ This serum-based therapy was formulated by challenging horses with inactivated infectious agents, which yielded potent antiserum that was then administered to diphtheria/tetanus patients. At the time, a rudimentary understanding of the molecular mechanisms underpinning the therapeutic efficacy of convalescent plasma complicated further development of immunotherapy against bacterial infections. Later, it came to be more clearly understood that antibodies targeting bacterial cells in the antiserum contributed directly to patient recovery.^2^ Opsonization of bacterial cells can promote their clearance *via* phagocytosis and/or complement.

In the years following von Behring’s seminal work, there was a large influx of small molecule antibiotics – an era ushered in by the discovery of penicillin. In a short period, small molecule antibiotics became the standard course of treatment against bacterial infections. More recently, traditional antibiotic therapy is becoming less effective as incidences of drug-resistant bacterial infections rise.^3, 4^ This trend can provide a powerful impetus to explore alternative strategies, including modern versions of the early work by von Behring that focuses on harnessing the patient’s own immune system for bacterial clearance.^5, 6^ The findings by von Behring serve as foundational precedence to the development of modern anti-bacterial immunotherapeutic agents, which can provide additional options for patients who are unresponsive to conventional antibiotics.

While modern immunotherapy has blossomed in the areas of oncology and autoimmunity, development of immunotherapies to treat bacterial infections has lagged behind. Nonetheless, increasing efforts towards this area have been made, as there are several monoclonal antibodies (mAbs) currently being developed against bacterial toxins^7–13^ and towards epitopes on the bacterial cell surface.^14–22^ One of the strategies being currently explored against cancer cells focuses on tagging cancer cell surfaces with small molecule haptens to promote their destruction by the immune system.^23–29^ For example, Low and coworkers have developed anti-cancer agents, which have undergone clinical evaluation, that are composed of folate linked to the endogenous hapten, dinitrophenol (DNP), to destroy cancer cells displaying high levels of folate receptors.^30–32^

Our group^33–37^ and others^38, 39^ have developed several classes of molecules that label the surface of bacterial pathogens with small molecule haptens. A principal design consideration for hapten display involves the choice of the targeting moiety to the bacteria cell surface. For Gram-positive pathogens, peptidoglycan (PG) is one of the most exposed epitopes on the cell surface. PG is a primary component of bacterial cell walls, and, in the case of Gram-positive bacteria, it can reach several nanometers in thickness. There are many structural components that are unique to PG including the inclusion of D-alanine (D-ala) residues within the stem peptide (humans do not biosynthesize D-ala^40, 41^). Herein, we describe the surface tagging of bacterial pathogens based on a peptide derived from a protein found in the innate immune system of *Ixodes scapularis* ticks called *Ixodes scapularis* antifreeze glycoprotein (IAFGP). It was recently found that IAFGP binds to the D-alanine residue within the PG of the bacterial cell wall.^42, 43^ Moreover, a fragment of the protein called P1 retained its ability to bind to the surface of the bacteria.^42, 44^ P1 has also been found to inhibit biofilm formation, potentiate antibiotics, and render mice resistant to septic shock.^42, 44, 45^ Given its therapeutic potential, we reasoned that P1 could additionally serve as a selective tagging modality to decorate the surface of bacterial cells. Here, we showed that conjugation of a non-native hapten resulted in high levels of antibody recruitment and immune-cell mediated uptake of bacterial cells (**Figure 1A**).

**Figure 1.**
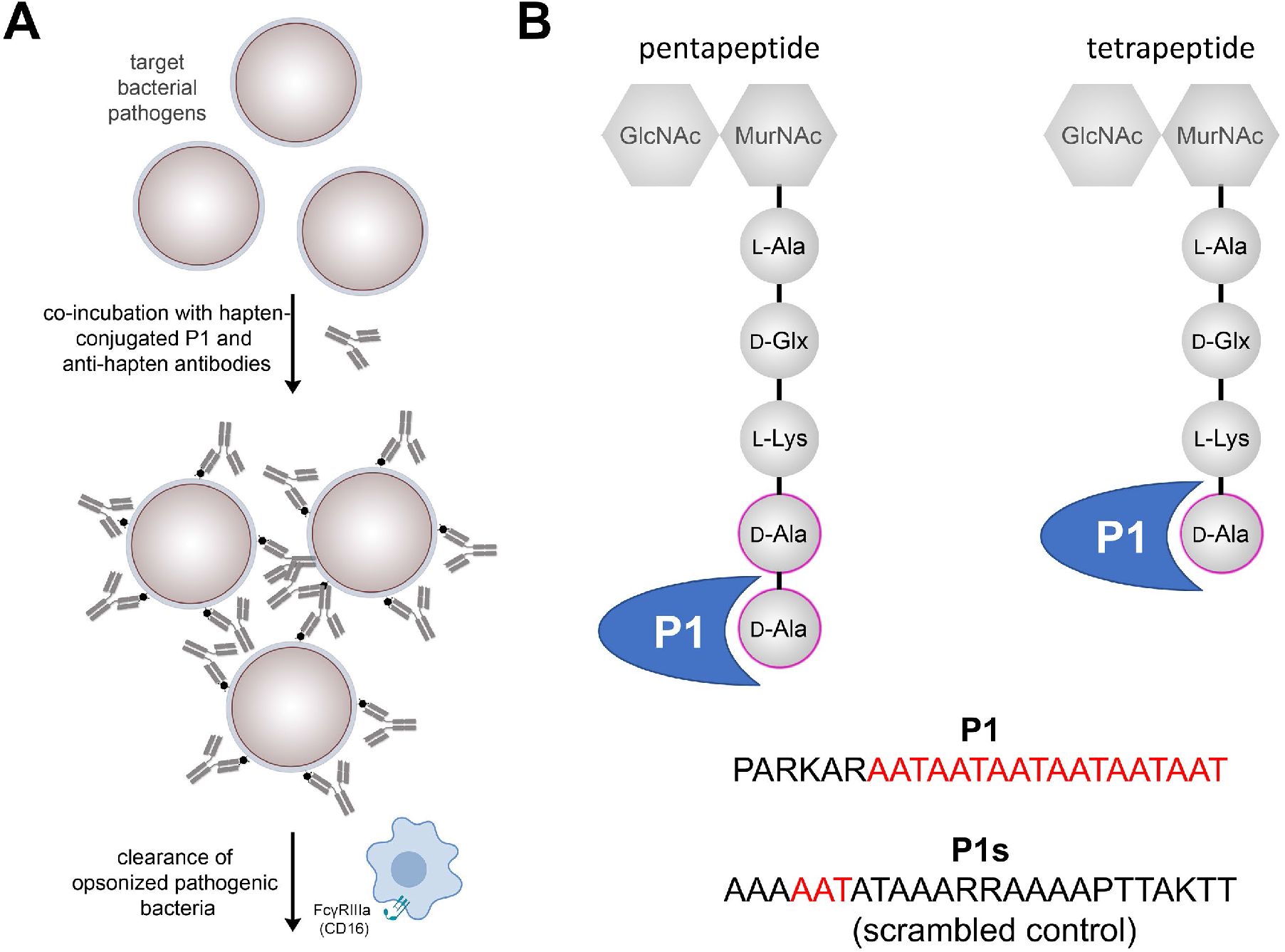
(**A**) General representation of hapten conjugates tagging the surface of bacteria followed by a specific immune response directed by the hapten. (**B**) Amino acid sequences of P1 and P1s with a cartoon representation of P1 binding to the D-Ala residue of a pentapeptide or tetrapeptide within the PG of the bacterial cell wall. D-iGlx refers to D-isoGln or D-isoGlu.

## Results and Discussion

Anti-bacterial immunotherapy has thus far been primarily based on the use of small molecule haptens for which humans already have an existing antibody pool (endogenous antibodies).^5, 46, 47^ Several different approaches by our group and others have demonstrated that using hapten-modified antibiotics, metabolic tags, and receptor binding polymers can harness this endogenous antibody pool to increase opsonization and subsequent killing of bacterial pathogens.^5, 35, 46–60^ However, despite such successes, there are shortcomings associated with endogenous haptens, such as a lack in temporal control of antibody circulation and variable levels of antibodies across a diverse population. Instead, we envisioned that a non-native hapten could overcome these challenges by the use of cognate exogenous mAbs. We describe a novel bifunctional molecule that associates with epitopes on the surface of pathogenic Gram-positive bacteria and displays an exogenous hapten for mAb recognition. This alternative approach can potentially reduce the necessity to discover and target pathogen-specific epitopes on the surface of Gram-positive bacteria. Targeting of bacterial cell surfaces was mediated by a peptide, P1, that binds to D-ala^42, 43^ present exclusively on the PG layers within the bacterial cell wall and not within the host (**Figure 1B**). P1 was labeled with a fluorescein moiety to promote the binding of exogenous anti-fluorescein antibodies to the surface of Gram-positive bacterial pathogens.

The cell wall of Gram-positive bacteria has a thick layer of PG on the extracellular side of the cytoplasmic membrane, whereas the PG of Gram-negative bacteria are found in the periplasmic space. The PG scaffold provides physical and chemical stability to the bacterial cell.^61, 62^ Generally, the structure of PG is highly conserved between species and is composed of repeating disaccharides connected to a short peptide, often referred to as the stem peptide. While there is variation in the stem peptide primary sequence, the canonical sequence is L-Ala-D-Glx-(L-Lys/m-DAP)-D-Ala-D-Ala where m-DAP is *meso*-diaminopimelic acid (**Figure 1B**). Often, proteins involved in the innate immune system will target the PG for mitigation of potential pathogens (e.g., lysozyme and Peptidoglycan Binding Proteins).^63, 64^ Similarly, the presence of the bacterial pathogen *Anaplasma phagocytophilum* in *Ixodes scapularis* ticks induces the host to express IAFGP.^42, 43^ Among the many biological activities related to IAFGP, it possess the ability to bind the terminal D-ala residue within the PG of the invading pathogen to alter bacterial cell permeability and interfere with biofilm formation.^42, 44^

IAFGP is unique in its repetitive nature of trimeric units (AAT) that are purported to bind D-Ala on bacterial PG.^65^ We envisioned that a fragment of IAFGP (peptide P1, PARKARAATAATAATAATAATAAT) could serve as a surface homing receptor to graft exogenous haptens onto the PG of the target bacteria (**Figure 2A**). To test the ability of the P1 peptide to bind to bacterial cell surfaces, P1 (and a scramble peptide P1s, AATAATATAAARRAAAAPTTAKTT) were synthesized using solid phase peptide synthesis (SPPS) and modified with a fluorescein on the *N*-terminus (**P1fl** and **P1fls**, respectively) (**Figure 2B**). Cell surface binding of **P1fl** and **P1fls** were evaluated against the Gram-positive bacterial species *Enterococcus faecalis (E. faecalis*) and *Enterococcus faecium (E. faecium*). We chose to investigate the binding of P1 to *E. faecalis* and *E. faecium* because both species can induce a drug resistant phenotype, thus making them difficult to treat with conventional antibiotics.^66, 67^ Stationary phase bacterial cells were incubated with **P1fl** and cellular fluorescence was analyzed by flow cytometry, whereby fluorescence levels are expected to correspond with binding of the bacterial cell surface (**Figure 2C, S1A**). Additionally, the PG scaffold within bacterial cells were metabolically labeled with diethylaminocumarin-modified D-amino acid, D-DADA, to easily identify overlapping fluorescence signals between bacterial cells and **P1fl** (**Figure S2, S1B**). Remarkably, high levels of surface labeling were observed for both *E. faecalis* and *E. faecium* when treated with **P1fl**, while treatment with **P1fls** did not result in bacterial surface tagging (**Figure 2C, S2C**). Moreover, we found that **P1fl** bound the surface of both *Enterococci* species in a concentration dependent manner (**Figure 2D, E**).

**Figure 2.**
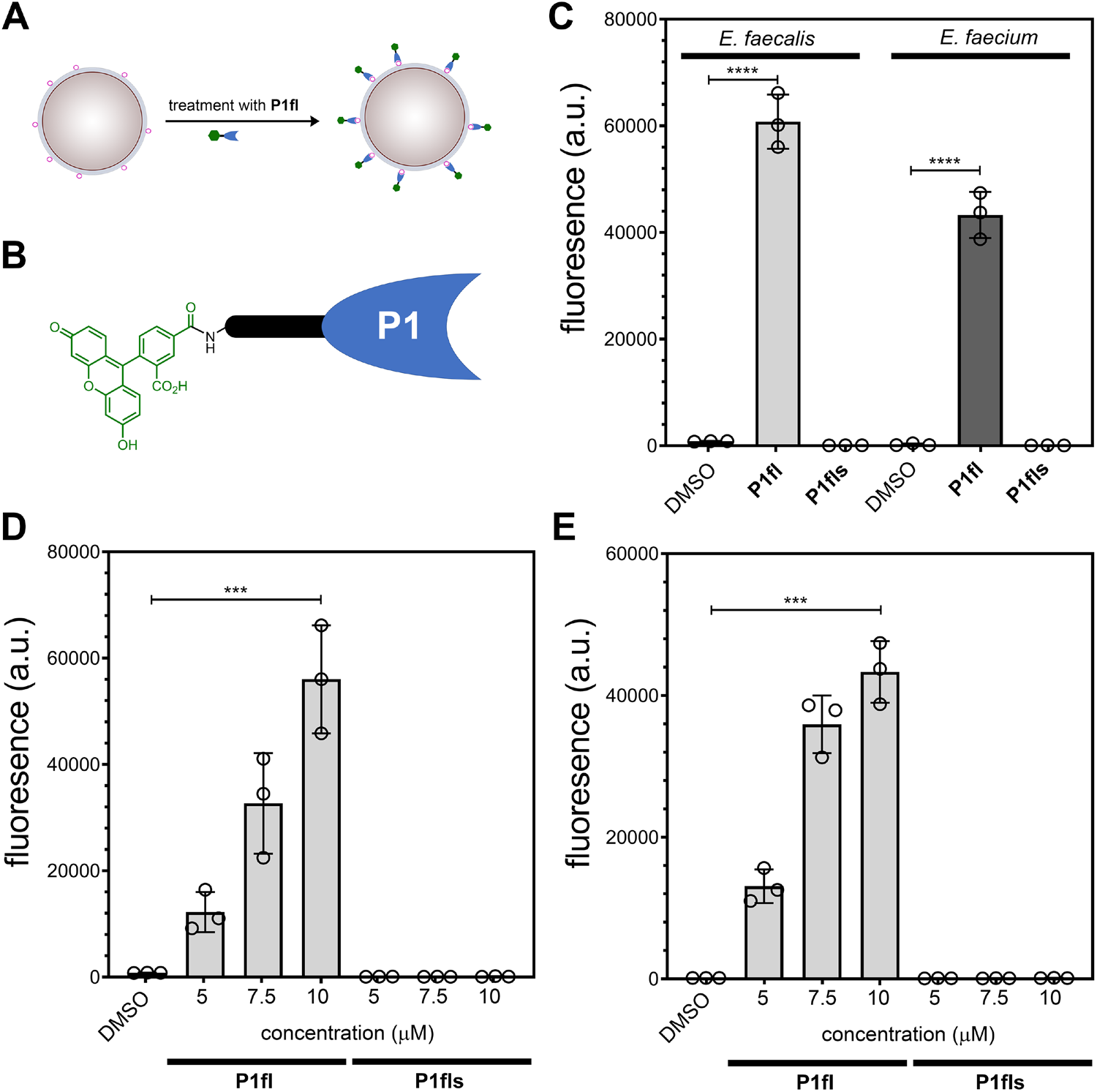
(**A**) Schematic of bacterial cells labeled with P1 probes. (**B**) Chemical structure of fluorescein modified to P1. (**C**) *E. Faecalis* 29212 and *E. faecium* 2127 were treated with 10 μM of P1 probes and analyzed by flow cytometry. (**C**) *E. Faecalis* 29212 and (**D**) *E. faecium* 2127 were treated with indicated concentrations of P1 probes and analyzed by flow cytometry. Data are represented as mean ± SD (n= 3). P-values were determined by a two-tailed t-test (*p < 0.05, **p < 0.01, ***p < 0.001, ****p < 0.0001, ns = not significant).

Further, we examined how **P1fl** binding would compare to vancomycin, given its well-established ability to bind to the D-ala-D-ala motif on lipid II or within the PG scaffold.^68, 69^ We anticipated that **P1fl** and vancomycin would exhibit similar cell labeling patterns being that both are expected to bind D-ala. *E. faecium* was co-incubated with BODIPY conjugated vancomycin (**VBD**)^70^ and tetramethyl-rhodamine modified P1 (**P1tam**) then analyzed by confocal microscopy (**Figure 3**). *E. faecium* displayed labeling of **VBD** at both the septal region and the surrounding cell wall as expected.^53, 71–73^ On the other hand, **P1tam** demonstrated labeling primarily at the surrounding cell wall, with minimal septal labeling. The septal region has the highest density of lipid II within the cell, as it is the site of PG biosynthesis; therefore, it is expected that vancomycin would display the highest degree of labeling to that region.^72^ The surrounding cell wall is primarily composed of enzymatically processed PG which contains mainly tetra- and tri- stem peptides, rather than penta- stem peptides.^74^ Moreover, we speculate that **P1tam** binding may be primarily directed to the surrounding cell wall because of its affinity for a singular D-ala motif rather than for D-ala-D-ala as with **VBD**. Additionally, this may be due in part to the limited accessibility of the septal region by P1, as it is larger than vancomycin.

**Figure 3.**
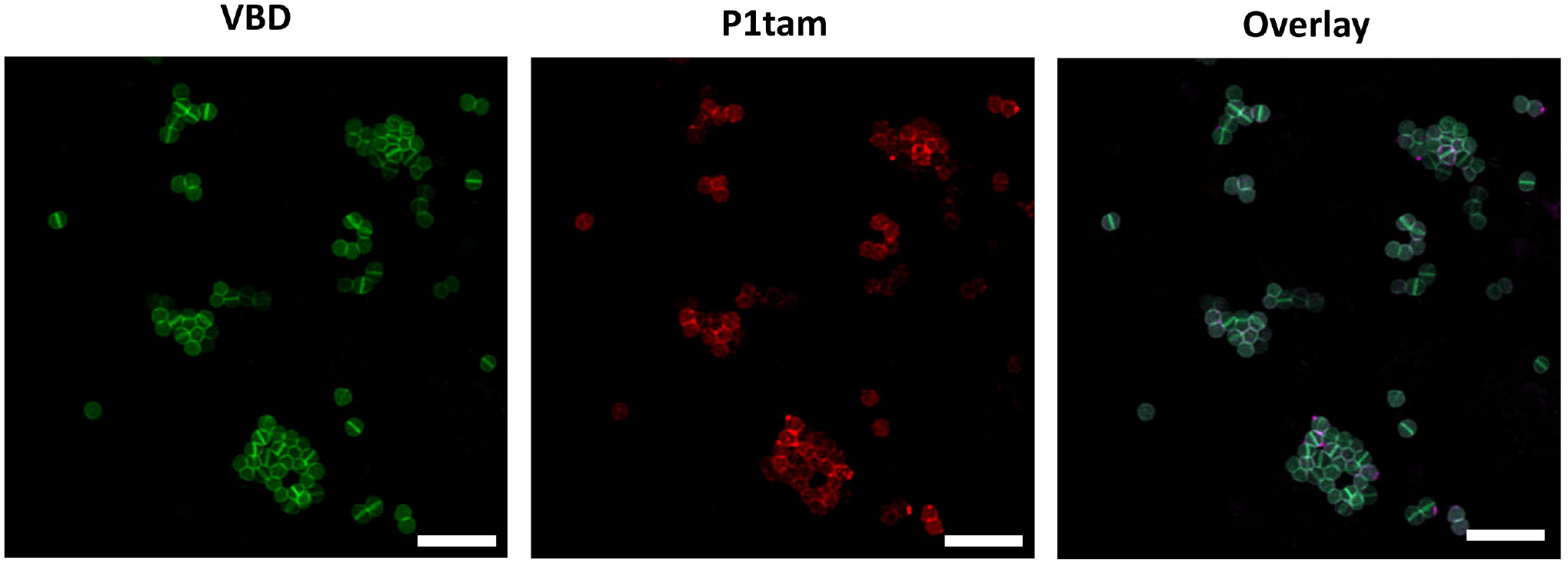
Confocal microscopy images of *E. faecium* 2127 treated with 10 μg/mL of VBD and 500 μM of P1tam. Shown is the overlay of channels corresponding to bodipy-Fl and tetramethyl-rhodamine. Scale bar = 3 μm.

Our lab has recently demonstrated a novel assay, SaccuFlow, to evaluate binding to bacterial PG using isolated, intact sacculi from a variety of bacterial strains (**Figure 4A**).^75^ To further demonstrate the binding specificity of **P1fl** to D-ala on PG, sacculus from *E. faecalis* and *E. faecium* isolated and individually incubated with **P1fl** and cell binding was assessed *via* flow cytometry (**Figure 4B**). As expected, **P1fl** bound to the sacculi of both *E. faecalis and E. faecium*, with *E. faecium* exhibiting higher binding levels as indicated by a higher fluorescence signal. We also identified that **P1fl** labeled both sacculi in a concentration dependent manner (**Figure S3A, B**). Additionally, incubation of **P1fls** exhibited background levels of fluorescence compared to **P1fl**, indicating that the scramble peptide did not bind to either strain of sacculi (**Figure 4B**). From these data, we were able to further demonstrate the binding propensity of **P1fl** to D-ala on PG *via* SaccuFlow and provide further evidence that binding indeed occurs with the PG of live bacterial cells.

**Figure 4.**
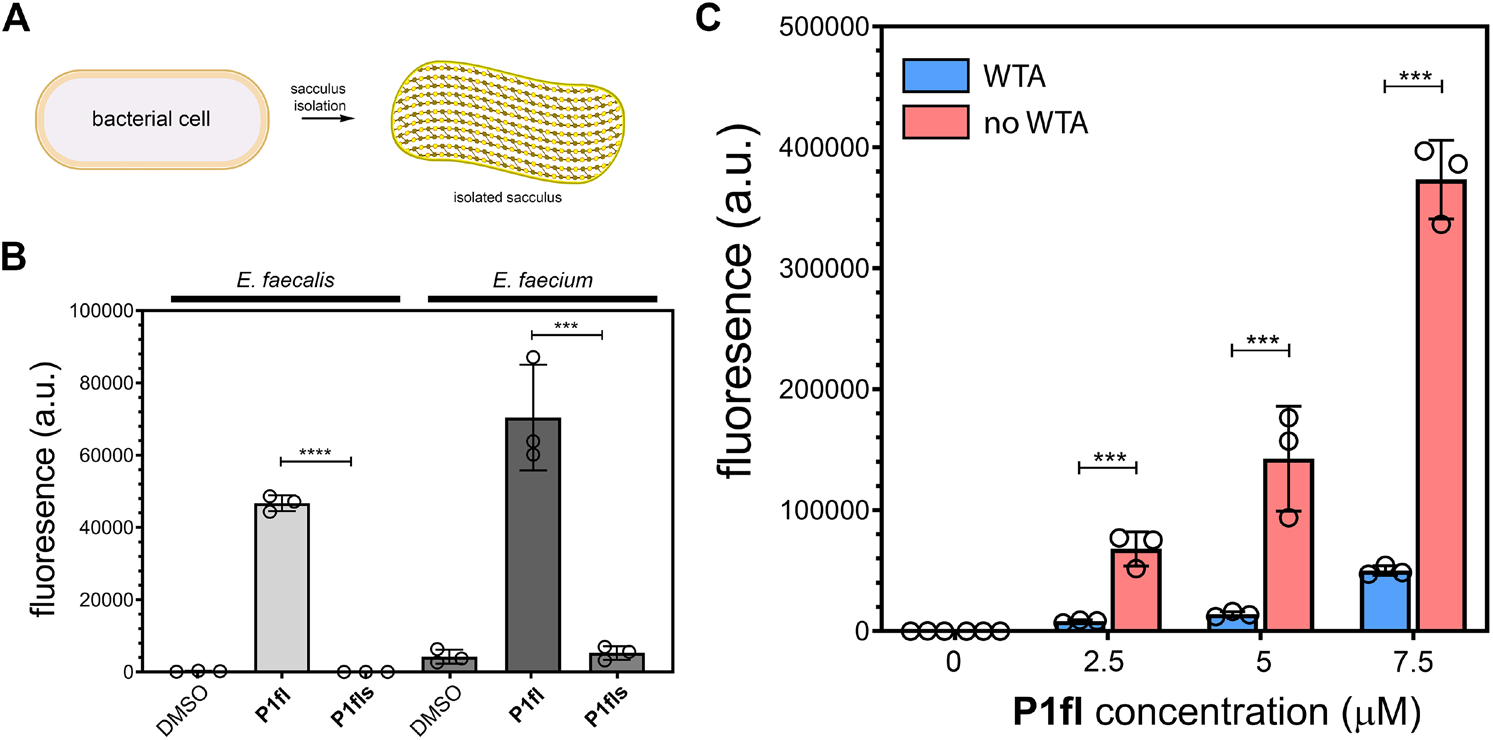
(**A**) Schematic of bacterial cells undergoing treatment to isolate intact bacterial sacculi. (**B**) *E. faecalis* 21922 and *E. faecium* 2127 sacculi were incubated with 7.5 μM of P1 probes and analyzed by flow cytometry. (**C**) *E. faecalis* 21922 sacculi with and without WTA were incubated with indicated concentration of **P1fl** and analyzed by flow cytometry. Data are represented as mean ± SD (n= 3). P-values were determined by a two-tailed t-test (*p < 0.05, **p < 0.01, ***p < 0.001, ****p < 0.0001, ns = not significant).

The accessibility of immune proteins to its bacterial cell wall target is largely mitigated by the presence of large surface polymers, wall teichoic acid (WTA) and lipoteichoic acid (LTA). Our lab,^76^ and others,^77–80^ have recently demonstrated that WTA play a major contribution to the accessibility of molecules to the PG of bacterial cells; therefore, we aimed to investigate the role that WTA plays on the accessibility of **P1fl** to the surface of *E. faecalis,* as WTA is the most highly abundant glycopolymer on the enterococci cell wall.^81, 82^ To achieve this, sacculi isolated from *E. faecalis* were treated with acid to hydrolyze the phosphodiester linkage between WTA and the PG, effectively removing the WTA from the isolated PG. Sacculi with and without WTA were incubated with **P1fl** and **P1fls**, and binding was monitored *via* flow cytometry. As expected, the removal of the WTA resulted in a large increase of cellular fluorescence when treated with **P1fl**, while binding of **P1fls** remained near background levels (**Figure 4C**). The binding of **P1fl** to the bacterial sacculi was observed in a concentration dependent manner for both with and without WTA and labeling was observed at concentrations as low as 2.5 μM.

Based on our findings that **P1fl** efficiently binds *Enterococci* bacterial cell surfaces, we anticipated that utilizing the fluorescein moiety as an exogenous hapten would result in opsonization of anti-FITC antibodies for a directed immune response to the bacterial cell. The directed opsonization of *E. faecalis* was anticipated to occur *via* two different pathways: a pre-assembly of **P1fl** with anti-FITC prior to bacterial cell binding and/or initial tagging of **P1fl** to the surface of *E.* f*aecalis* followed by the sequential binding of anti-FITC (**Figure 5A**). We opted to maintain the fluorescein hapten as it has been shown to successfully direct the recruitment of anti-FITC antibodies for killing of cancer cells.^30, 83^ Additionally, to improve surface presentation of the hapten to anti-fluorescein antibodies, we positioned the fluorescein moiety to be displayed on the γ amine of an additional *N*-terminal lysine residue on P1 (**Figure 5B**). We directed our efforts of antibody recruitment towards *E. faecalis*, as **P1fl** demonstrated high levels of cell binding and has vital clinical applicability, being that some types of drug resistant *E. faecalis* can induce a vancomycin-resistant phenotype.^84, 85^ Vancomycin-resistant *Enterococci* (VRE) are identified as serious threat level pathogens by the Centers for Disease Control and Prevention (CDC), as treatment options for these organisms are becoming limited.^86, 87^ Vancomycin-sensitive (VSE) *E. faecalis* was incubated with a solution containing **P1fl** or **P1Kfl** and AlexaFluor 647 anti-FITC antibodies, and then analyzed by flow cytometry, where fluorescence levels is expected to correspond with binding of anti-FITC antibodies (**Figure 5C**). ^88, 89^ VSE *E. faecalis* demonstrated greater levels of antibody recruitment when treated with **P1Kfl** at as low as 5 μM compared to **P1fl,** despite **P1fl** exhibiting much higher cell labeling levels over **P1Kfl** (**Figure S4**). This confirms that hapten presentation to the antibody plays a pivotal role in antibody recognition.

**Figure 5.**
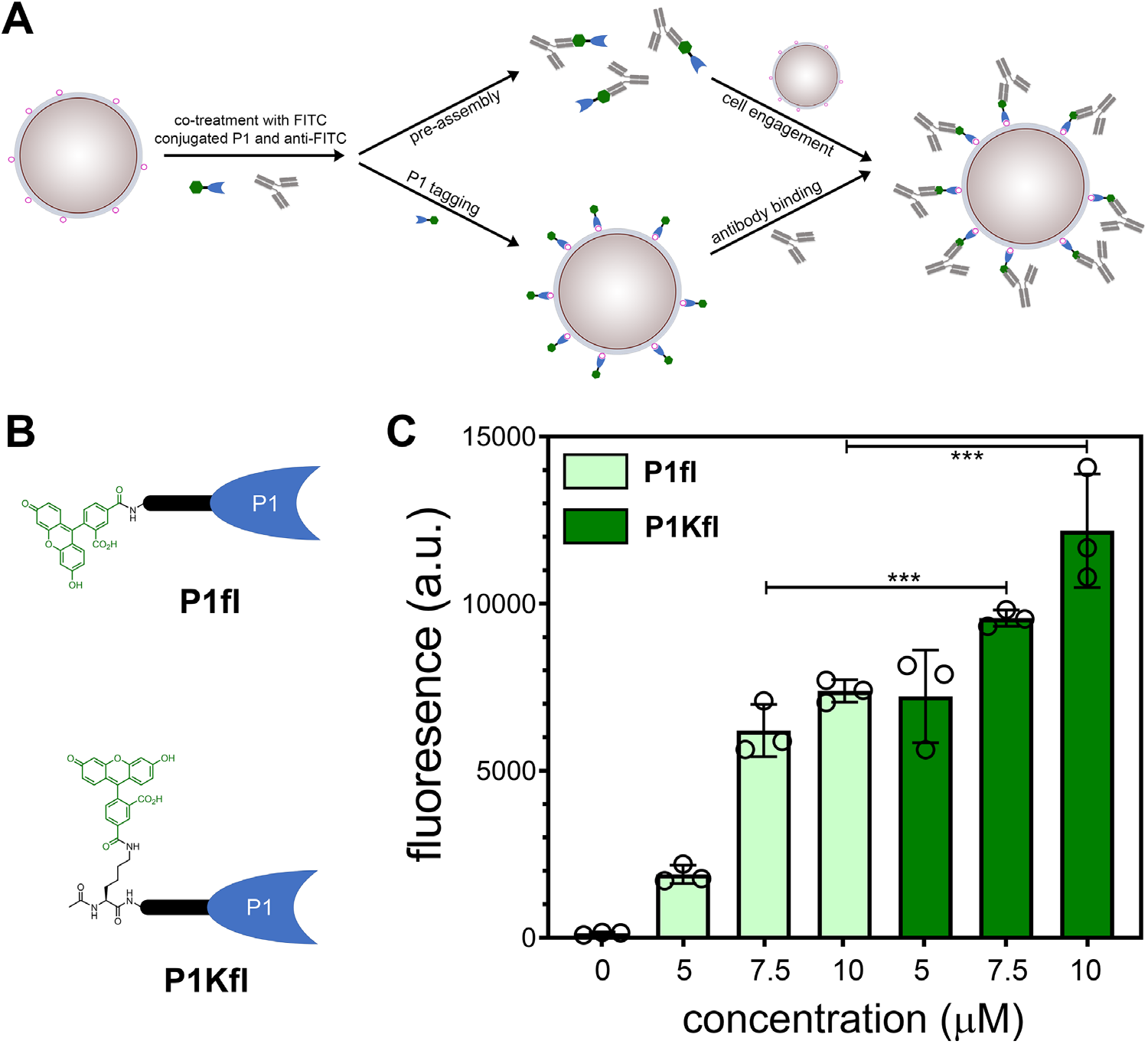
(**A**) Cartoon representation of co-incubation of bacterial cells with P1 probes and anti-FITC antibodies. Cell targeting can occur via two pathways: pre-assembly of P1fl/anti-FITC prior to cell engagement or P1 tagging followed by antibody binding. (**B**) Chemical structures of modifications to P1 peptides. (**C**) *E. faecalis* 21922 was treated with indicated concentrations of P1 probes and AlexaFluor 647 anti-FITC antibodies where anti-FITC fluorescence was measured by flow cytometry. Data are represented as mean ± SD (*n*= 3). *P*-values were determined by a two-tailed *t*-test (\**p* < 0.05, \*\**p* < 0.01, \*\*\**p* < 0.001, ns = not significant).

Next, we wanted to evaluate the cell labeling and antibody recruitment of VSE *E. faecalis* compared to vancomycin-resistant (VRE) *E. faecalis*. When VSE and VRE *E. faecalis* were treated with **P1Kfl** immune tags, VRE *E. faecalis* exhibited much higher levels of cell labeling compared to VSE, as indicated by a higher fluorescence signal (**Figure 6A**). The same trend in fluorescence was observed for the recruitment of anti-FlTC antibodies, whereby **P1Kfl** treatment of VRE *E. faecalis* led to much higher levels of fluorescence relative to VSE (**Figure 6B**). **P1Kfl** also demonstrated superior levels of antibody recruitment to VRE *E. faecalis* compared to when treated with **P1fl**. Further, background fluorescence levels were observed for all scramble P1 peptides for both cell labeling and antibody recruitment in VRE *E. faecalis* (**Figure 6A, B**). We speculate that the differences in recruitment between VSE and VRE *Enterococci* may be reflective of several variations in surface composition of drug resistant phenotypes including chages to cell surface accessibily. It may be possible that varying amounts of LTA and WTA of VSE and VRE may contribute to **P1Kfl** binding and subsequent anti-FITC recruitment.^81, 82^

**Figure 6.**
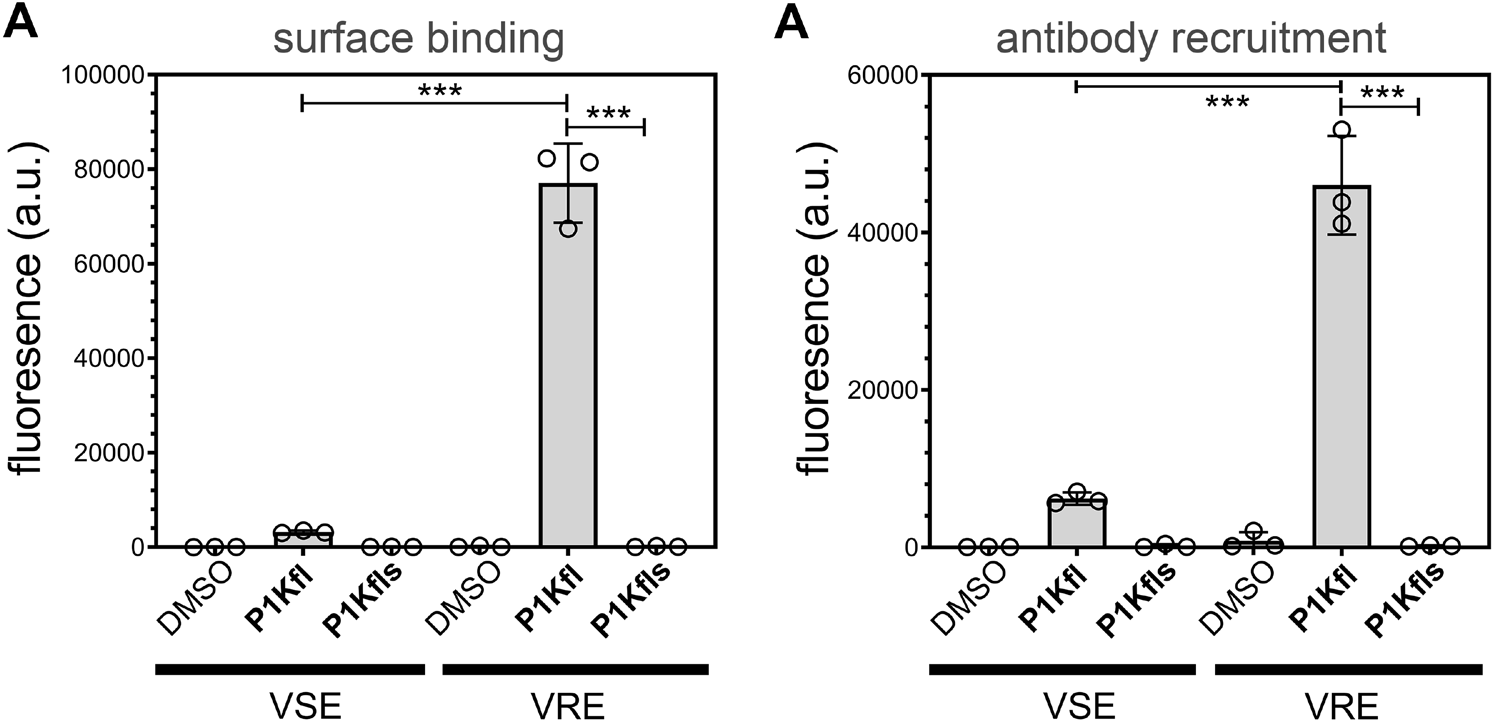
(**A**) *E. faecalis* 21922 (vancomycin sensitive, VSE) and *E. faecalis* 51922 (vancomycin resistant, VRE) were treated with 7.5 μM of P1 probes and FITC fluorescence was measured by flow cytometry. (**B**) VSE and VRE *E. faecalis* were treated with 7.5 μM of P1 probes and AlexaFluor 647 anti-FITC antibodies and anti-FITC fluorescence was measured by flow cytometry. Data are represented as mean ± SD (n= 3). P-values were determined by a two-tailed *t*-test (*p < 0.05, **p < 0.01, ***p < 0.001, ns = not significant).

It has been demonstrated that in order to induce the drug resistant phenotype of *E. faecalis*, vancomycin must be supplemented to the growth medium.^90–92^ We set out to examine how the addition of vancomycin to the growth medium would affect binding of **P1Kfl** and subsequent anti-FITC recruitment. Interestingly, minimal differences in bacterial cell binding and antibody recruitment were observed between induced and non-induced VRE *E. faecalis* (**Figure S5A, B**). We speculate that the ability of the P1 peptide to bind terminal D-ala, rather than D-ala-D-ala as with vancomycin, would remain unchanged following the induction of the VRE because this would not be expected to change the overall levels of D-ala. Critically, our results show that P1 conjugates would likely not be impacted by this type of VRE drug resistant phenotype.

The specificity of anti-FlTC for **P1Kfl** was further confirmed by treating cells with a mock antibody in place of anti-FITC; background fluorescence levels were observed for both P1 and P1s probes in the presence of Alexa Fluor 647 conjugated anti-human IgG (**Figure S6A**). Additionally, anti-FITC recruitment was not disrupted by the presence of pooled human serum, indicating that the **P1Kfl**/anti-FITC complex maintains recognition of the bacterial cell surface in a complex medium (**Figure S6B**). Based on our findings, we can conclude that fluorescein was grafted on the surface of VRE and successfully opsonized with anti-FITC antibodies.

To further expand the applicability of the P1 peptide to graft haptens on to the surface of bacterial cells, and show its overall generality in surface tagging, an endogenous hapten, dinitrophenol (DNP), was used in place of the FITC moiety on P1, resulting in **P1dnp** (**Figure S7A**). Anti-DNP antibodies make up 1-2% of the naturally occurring human antibody pool.^66, 67, 93^ Our group^33–36^ and others^38, 39^ have demonstrated that DNP can be used to recruit anti-DNP antibodies from human serum for bacterial immunotherapy. As expected, when **P1dnp** and anti-DNP were incubated with VRE *E. faecalis,* high fluorescence levels were only observed with **P1dnp** treatment and not with the scrambled sequence **P1dnps** (**Figure S7B**). Based on these results, a hapten modified P1 can function as a surface tagging modality for the recruitment of both exogenous and endogenous antibodies to pathogenic bacteria.

Finally, we set out to determine whether anti-FITC antibodies present on the surface of VRE *E. faecalis* would increase recognition and phagocytosis of the bacterial cells by macrophages, as antibody dependent cellular phagocytosis (ADCP) is one of the primary mechanisms for clearance of bacterial pathogens by the immune system.^94^ Bacterial cells were co-incubated with J774A macrophages in the presence of **P1Kfl** and anti-FITC antibodies, with phagocytosis of bacterial cells being tracked by flow cytometry. VRE *E. faecalis* uptake by macrophages increased by 20-fold in the presence of both **P1Kfl** and anti-Fl antibodies, compared to macrophages co-incubated with VRE *E. faecalis* treated with **P1Kfls**. Background levels of phagocytosis were observed when anti-FITC antibodies were omitted for both **P1Kfl** and **P1Kfls** (**Figure 7A, B**). Confocal microscopy confirmed that the fluorescent population of bacterial cells were phagocytosed into the macrophages (**Figure 7C**). This demonstrates that increasing antibody opsonization of VRE can enhance phagocytosis and clearance of bacterial pathogens.

**Figure 7.**
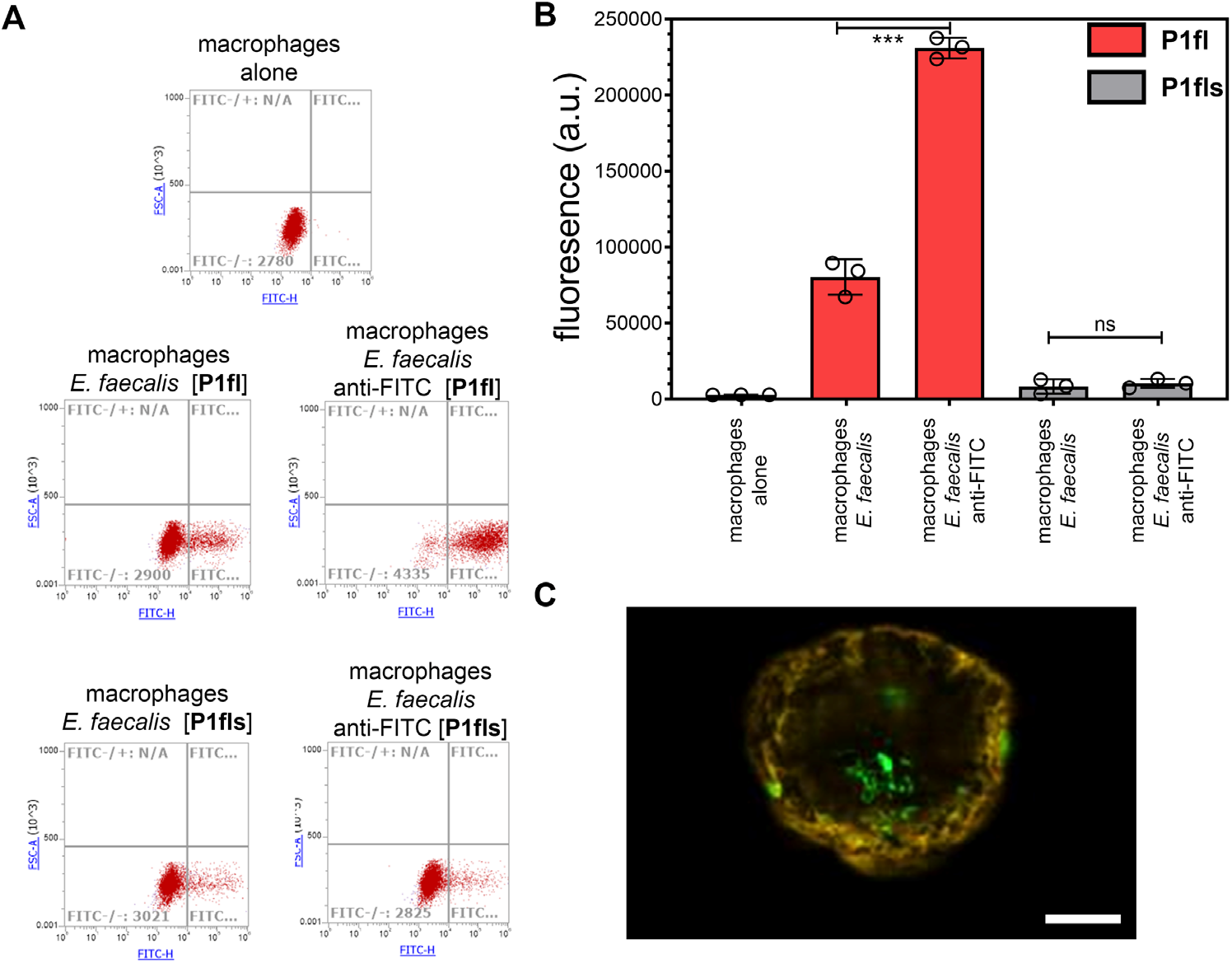
(**A**) Representative scatter plots exhibiting changes in fluorescence for the macrophage population co-incubated with P1 probe opsonized *E. faecalis* 51922. (**B**) Flow cytometry quantification of *E. faecalis* cells treated with P1 probes and anti-FITC antibodies prior to co-incubation with J774A macrophages. Data are represented as mean ± SD (n= 3). P-values were determined by a two-tailed t-test (*p < 0.05, **p < 0.01, ***p < 0.001, ns = not significant). (**C**) Confocal microscopy images of *E. faecalis* treated with 7.5 μM **P1fl** and anti-FITC phagocytosed into macrophages. Cells were fixed with formaldehyde and treated with tetramethyl-rhodamine-tagged WGA (5 μg/mL) for 30 min. Shown is the overlay of the channels corresponding to FITC and tetramethyl-rhodamine. Scale bar = 10 μm.

## Conclusion

In conclusion, we have shown that **P1fl** is able to bind to whole bacterial cell surfaces in a sequence dependent manner, as a scrambled version of P1 demonstrated no binding propensity towards bacterial cells. We were able to label the surface of two strains of *Enterococci*, *E. faecalis* and *E. faecium*, using **P1fl** as demonstrated by flow cytometry and confocal microscopy. Additionally, **P1Kfl** was able to induce the recruitment of exogenous anti-FITC antibodies to the surface of *E. faecalis* and facilitated a P1-mediated immune cell uptake of the bacterial cells. We demonstrated that the hapten displayed on P1 can be altered to recruit its cognate antibodies, thus demonstrating the broad applicability of P1 to target bacterial cell surfaces with various treatment modalities. Additionally, **P1fl** preferentially targeted the PG of vancomycin resistant *E. faecalis*; therefore, **P1fl** may be used as a therapeutic option in conjunction with antibiotics for the treatment of drug resistant *Enterococci* species.

## Acknowledgments

B.S. and M.M.P. were supported by the NIH grant GM124893-01.

