## Supporting Information for "Immuno-targeting of Gram-positive Pathogens *via* a Cell Wall Binding Tick Antifreeze Protein"

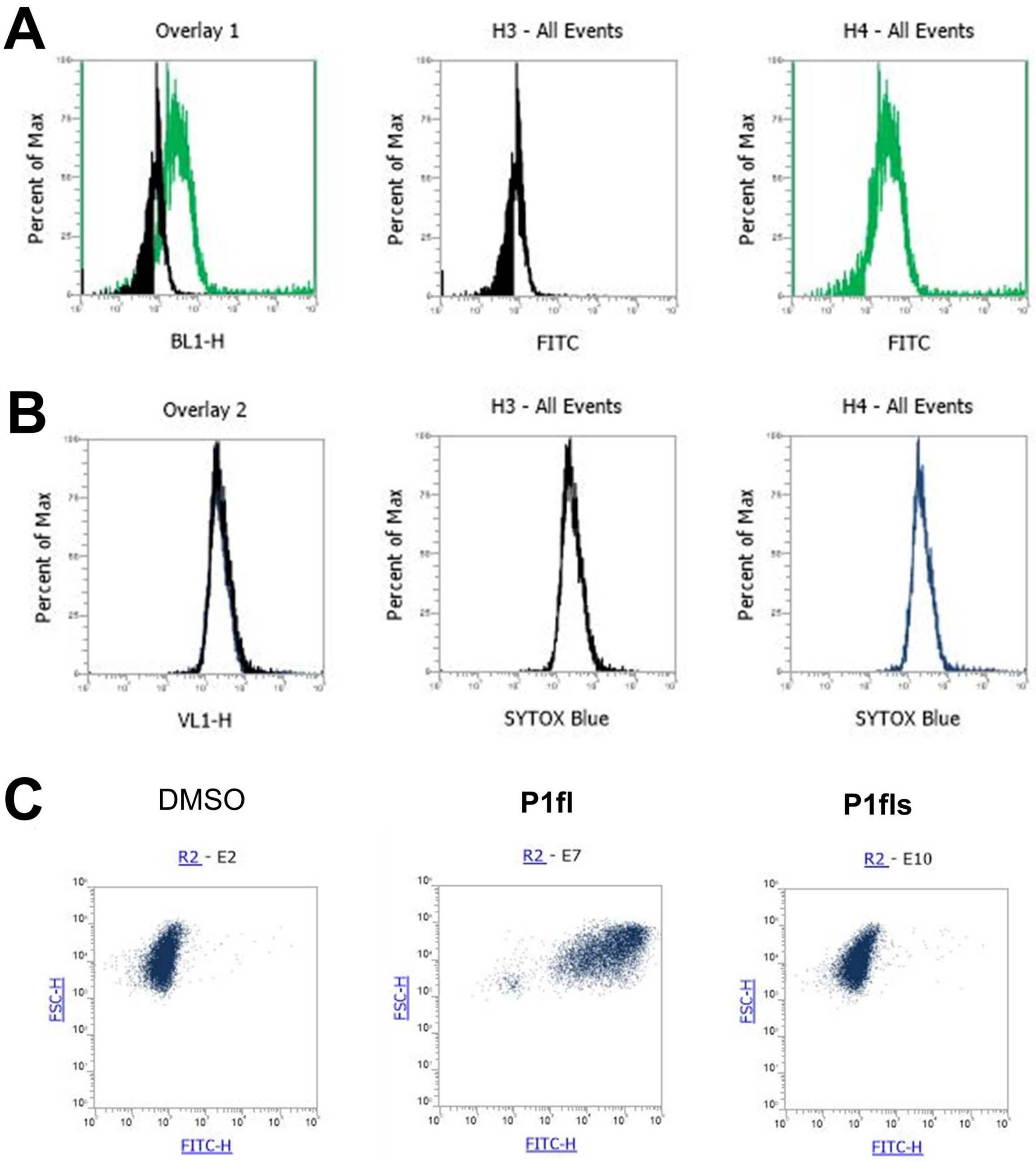


**Supplemental Figure S1.** (A) Flow cytometry histogram plot representing a shift in fluorescence between unlabeled *E. faecalis* 29212 cells (black) and cells treated with P1fl (green). (B) Flow cytometry histogram plot representing the fluorescence signal for DADA treated *E. faecalis* cells also treated without (black) and with P1fl (blue). (C) Representative flow cytometry dot plot indicating changes in fluorescence of the *E. faecium* cell population when incubated with P1 probes.

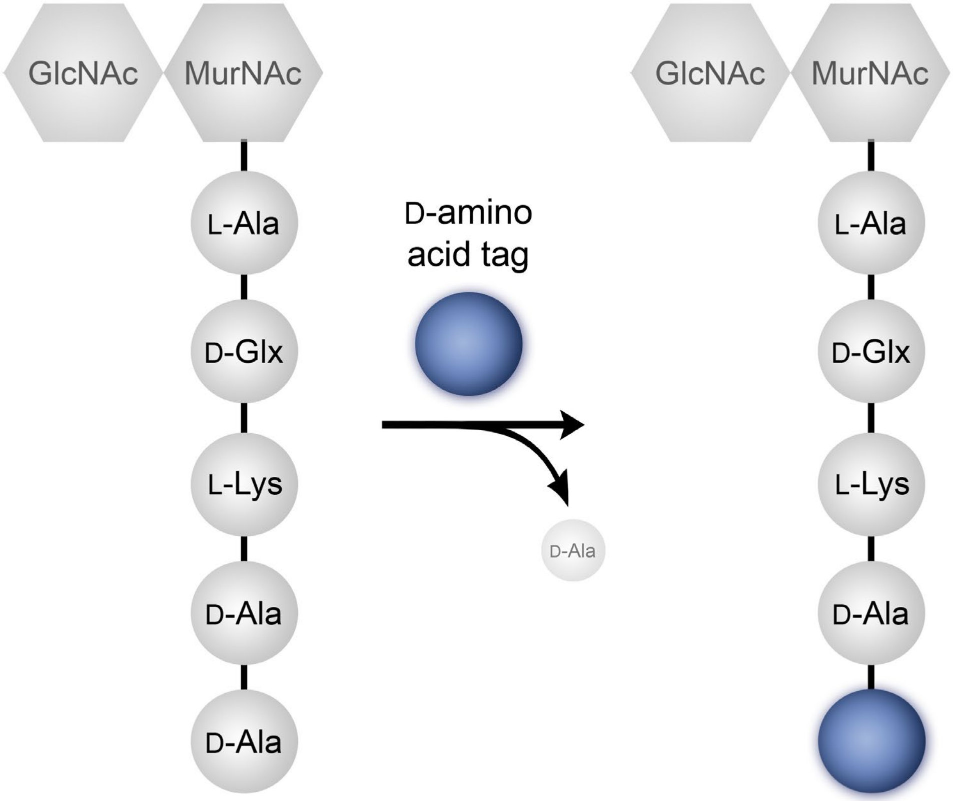


**Supplemental Figure S2.** Mode of incorporation of single amino acids into the 5th position of the peptidoglycan stem peptide.

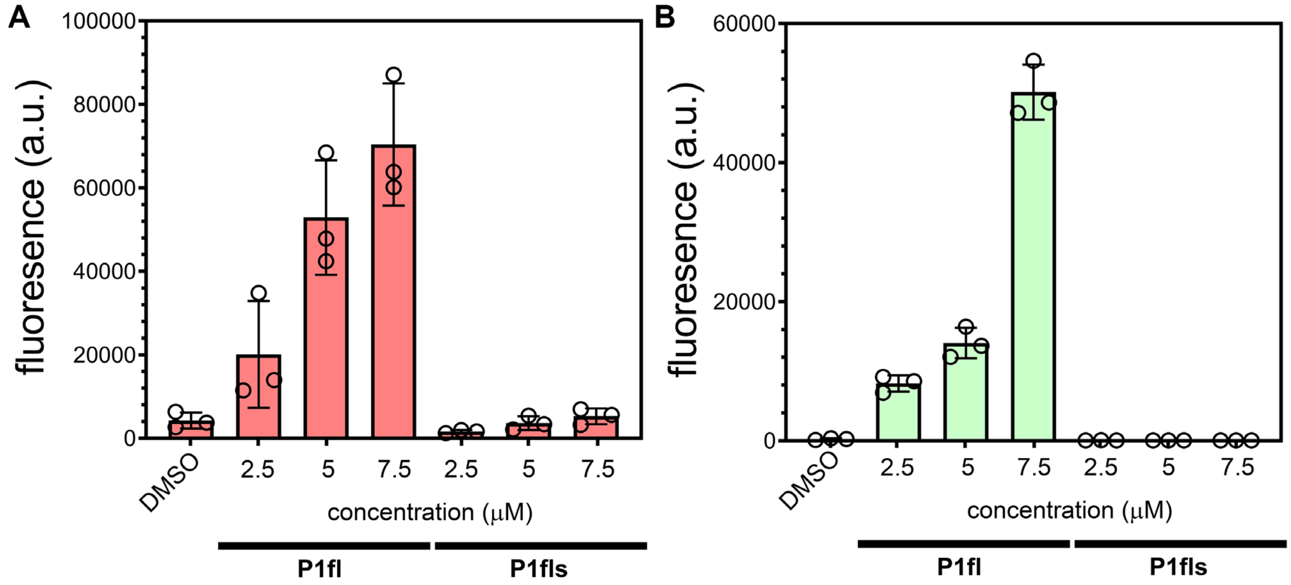


**Supplemental Figure S1.** (**A**) *E. faecium* 2127 sacculi and (**B**) *E. faecalis* 29212 sacculi were treated with indicated concentrations of P1 probes and analyzed via flow cytometry. Data are represented as mean ± SD (n= 3). P-values were determined by a two-tailed t-test (*p < 0.05, **p < 0.01, ***p < 0.001, ns = not significant).

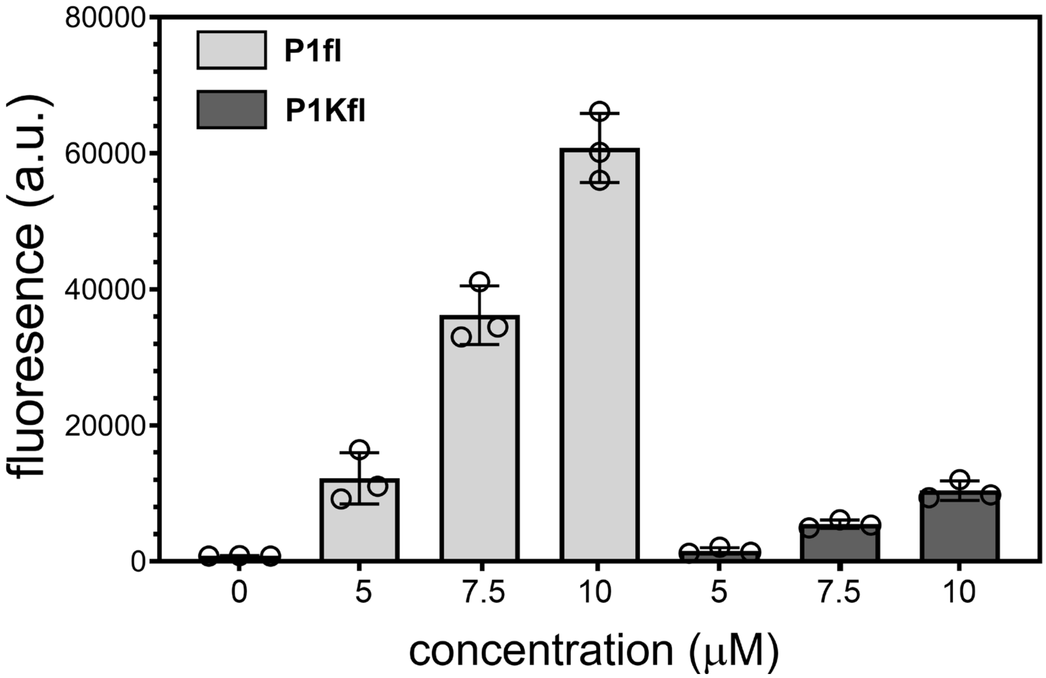


**Supplemental Figure S2.** *E. faecalis* 29212 was treated with indicated concentrations of P1 probes and analyzed for cell binding. FITC fluorescence was measured by flow cytometry. Data are represented as mean ± SD (n= 3). P-values were determined by a two-tailed t-test (*p < 0.05, **p < 0.01, ***p < 0.001, ns = not significant).

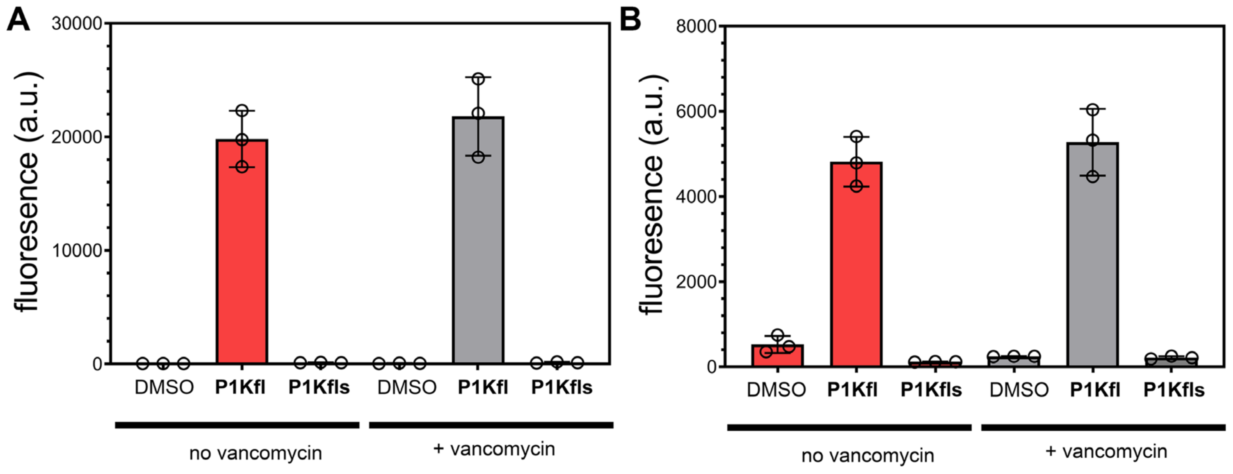


**Supplemental Figure S3.** *E. faecalis* 51922 cells grown with or without 16 ug/mL of vancomycin and analyzed for (**A**) cell binding and (**B**) anti-FITC recruitment when treated with P1 probes. Samples were analyzed via flow cytometry. Data are represented as mean ± SD (n= 3). P-values were determined by a two-tailed t-test (*p < 0.05, **p < 0.01, ***p < 0.001, ns = not significant).

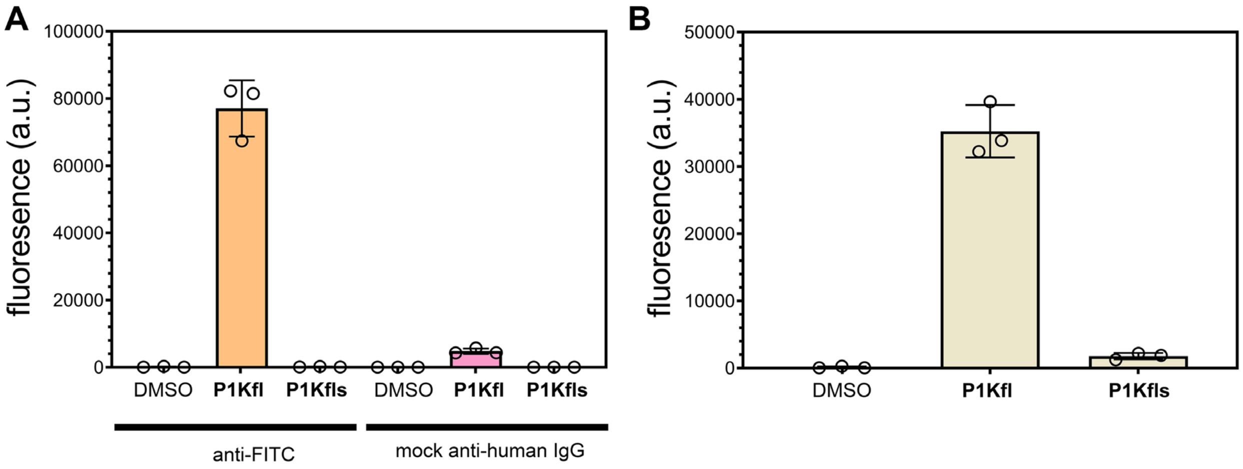


**Supplemental Figure S4.** Flow cytometry data for *E. faecalis* 51922 cells treated with (A) P1 probes and either anti-FITC or anti-human IgG (mock antibody) and (B) P1 probes + anti-FITC in the presence of human serum. Data are represented as mean ± SD (n= 3). P-values were determined by a two-tailed t-test (*$p < 0.05$, **$p < 0.01$, ***$p < 0.001$, ns = not significant).

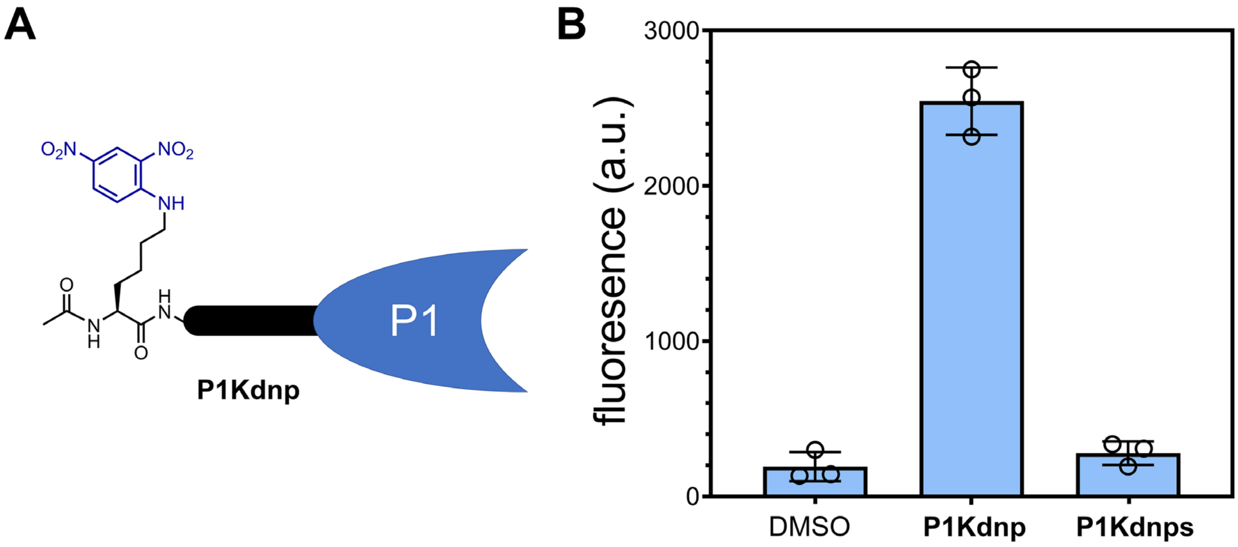


**Supplemental Figure S5**. (A) Cartoon representation of P1 probe modified with a DNP hapten. (B) Flow cytometry analysis of *E. faecalis* 51299 cells treated with 10 μM of P1 probes. Data are represented as mean ± SD (n= 3). P-values were determined by a two-tailed t-test (*p < 0.05, **p < 0.01, ***p < 0.001, ns = not significant).

**Materials.**

All peptide related reagents and protected amino acids were purchased from Chem-Impex. 5, 6-carboxyfluorescein was purchased from Chem-Impex. Anti-fluorescein and anti-human IgG was purchased from Jackson Immuno Research Laboratories. Anti-dinitrophenol KHL was purchased from Thermo Fisher. Pooled Human Serum was purchase from Sigma Aldrich. Dulbecco's Modified Eagle's Medium (DMEM) was purchased from VWR. Fetal Bovine Serum (FBS) was purchased from R&D Systems. Penicillin-Streptomycin was purchased from Sigma-Aldrich. All other organic chemical reagents were purchased from Fisher Scientific or Sigma Aldrich and used without further purification.

**Methods.**

**Bacterial Cell Culture.** Bacterial cells were cultured in specified media in an aerobic environment while shaking at 250 rpm at 37 °C. *E. faecalis* 29212 (vancomycin- sensitive) and *E. faecalis* 51299 (vancomycin- resistant) were grown in brain heart infusion broth (BHI). *E. faecium* ATCC BAA 2127 (vancoymycin-sensitive) and *E. faecium* ATCC BAA 2317 (vancoymycin-resistant) were grown in Trypticase Soy Broth (TSB). *E. faecalis* 51299 and *E. faccium* 2317 were supplemented with 16 µg/mL vancomycin to induce resistant phenotype. BLS2 organisms should be manipulated using proper protective equipment.

**Mammalian Cell Culture.** J774A.1 cells were cultured in Dulbecco's modified Eagle's medium (DMEM) supplemented with 10% (v/v) FBS, 50 IU/mL penicillin, 50 ug/mL streptomycin, and 2 mM L-glutamine in a humidified atmosphere of 55 $CO_2$ at 37 $^oC$.

**Fluorescent Labeling of Whole Bacterial Cells with P1 conjugates.** Bacterial cells were grown over-night to stationary phase in corresponding growth media while shaking (250 rpm) at 37 $^oC$. Bacterial cells from the overnight growth were used to inoculate BHI (1:100) supplemented with 100 µM DADA and incubated at 37 °C with shaking (250 rpm) for 16 h. The bacteria were harvested, washed three times with 1X phosphate buffered saline (PBS). The bacterial cells were resuspended in 1X PBS supplemented with P1- or P1s- fl conjugates (7.5 µM or designated concentration) and incubated at 37$^o$C with shaking (250 rpm) for 30 min. The bacteria were washed two times with 1X PBS, fixed with 2% formaldehyde solution, and analyzed using the Attune NxT Flow Cytometer (Thermo Fischer) equipped with a 405 nm and a 488 nm laser with 440/50 and 530/30 nm bandpass filter respectively. The data were analyzed using Attune Nxt software.

**Confocal Microscopy Analysis.** Bacterial cells were grown over-night to stationary phase in corresponding growth media while shaking (250 rpm) at 37 $^oC$. Bacterial cells from the over-night growth were harvested and washed once with 1X PBS. The bacterial cells were resuspended in 1X PBS supplemented with 500 µM of P1tam and 10 µg/mL of vancomycin bodipy Fl conjugate (Invitrogren #V34850, VBD) and incubated at 37$^o$C with shaking (250 rpm) for 30 min. The bacteria were washed three times with 1X PBS, fixed with 2% formaldehyde solution, and imaged using the Zeiss LSM 980 Imaging

System with multiplex Airyscan. We acknowledge the Keck Center for Cellular Imaging for the usage of the Zeiss 880/980 multiphoton Airyscan microscopy system (PI- AP: NIH-OD025156).

**Peptidoglycan Isolation of *E. faecalis*.** *E. faecalis* 29212 (vancomycin sensitive) and *E. faecalis* 51299 (vancomycin resistant) were grown over-night to stationary phase in BHI medium. A 100 mL culture volume in BHI medium was inoculated (1:100) from the stationary phase cultures and allowed to grow for 16 h while shaking (250 rpm) 37 $^{o}$C. The cultures were harvested, resuspended in 1X PBS, boiled at 100 $^{o}$C for 25 min, and centrifuged at 14,000 g for 15 min at 4 $^{o}$C. The cells were placed in 50 mL of 2% (w/v) sodium dodecyl sulfate (SDS) and boiled for 30 min followed by centrifugation at 14,000 g for 15 min at 4 $^{o}$C. Cells were then washed 6 times DI water to remove all the SDS. After washing, cells were resuspended in 25 mL of 20 mM Tris buffer (pH 8.0). Pellets were treated with 800 μg DNase for 24 hours followed by 800 μg trypsin for another 24 h at 37 $^{o}$C while shaking (115 rpm). Pellets were boiled for 25 min followed by centrifugation at 14,000 g for 15 min at 4 $^{o}$C. Pellets were resuspended in 1M HCl for 4 h at 37 $^{o}$C with shaking to remove wall teichoic acids. The pellet was harvested by centrifugation at 4,000 rpm for 10 min and washed with DI water until the pH of the supernatant reached 5-6. The final pellet was resuspended in 1X PBS, and further diluted for analysis by flow cytometry.

**Peptidoglycan Isolation of *E. faecium*.** *E. faecium* BAA 2127 (vancomycin sensitive) and *E. faecium* BAA 2317 (vancomycin resistant) were grown over-night to stationary phase in TSB medium. The same protocol was followed as stated prior for peptidoglycan isolation of *E. faecalis*.

**Fluorescent Labeling of Bacterial Sacculi.** Bacterial sacculi were resuspended in 1X PBS supplemented with P1fl or P1fls (at designated concentration) and incubated at 37$^{o}$C with shaking (250 rpm) for 30 min. Samples were washed twice with 1X PBS, fixed in a 2% formaldehyde solution, and analyzed by flow cytometry as previously described.

**Co-incubation of Bacterial Cells with P1 conjugates and anti-FITC antibodies.** Bacterial cells were grown over-night to stationary phase in corresponding growth media while shaking (250 rpm) at 37 $^{o}$C. Bacterial cells from the overnight growth were used to inoculate BHI (1:100) supplemented with 100 µM DADA and incubated at 37 °C with shaking (250 rpm) for 16 h. The bacteria were harvested, washed three times with 1X PBS. The bacteria were resuspended in 50 μL of PBS containing P1fl and P1fls (7.5 μM or indicated concentration), 10 % (v/v) FBS, and 0.03 μg/mL Alexa Fluor 647-conjugated mouse anti-fluorescein (FITC) (Jackson Immuno Research #200-602-037) and incubated at 37$^{o}$C for 30 min with shaking (250 rpm) protected from light. Samples were washed twice with 1X PBS, fixed in a 2% formaldehyde solution, and analyzed by flow cytometry (as described above) equipped with a 637 nm laser with 670/14 nm bandpass filter.

**Co-incubation of Bacterial Cells with P1 conjugates and anti-DNP antibodies.** Bacterial cells were prepared as described prior, with the exception that the bacteria were resuspended in 50 μL of 1X PBS containing P1dnp or P1dnps (10 μM or indicated concentration), 10 % (v/v) FBS, and 0.04 μg/mL Alexa Fluor 488-conjugated rabbit anti-dinitrophenol KHL (Thermo Fischer #A-11097) incubated at 37°C for 30 min with shaking (250 rpm) protected from light. Samples were washed twice with 1X PBS, fixed in a 2% formaldehyde solution, and analyzed using the Attune NxT Flow Cytometer (Thermo Fischer) equipped with a 488 nm laser and 525/40 nm bandpass filter. The data were analyzed using Attune Nxt software.

**Incubation of bacterial cells with P1 conjugates and anti-FITC antibodies in human serum.** Pooled human serum (PHS, Sigma-Aldrich #H4522) was diluted to 25% in 1x PBS and incubated with bentonite for 20 min at 37 °C to deactivate the lysozyme. The serum supernatant was obtained and diluted to a final concentration of 10% for use in the assay. The bacterial cell labeling and antibody recruitment protocol was followed as previously described, except for the 10% PHS was used rather than 10% FBS.

**Incubation of bacterial cells with P1 conjugates and a mock antibody.** The same protocol was followed as stated prior but with Alexa Fluor 647 goat anti-human IgG (Jackson Immuno Research #109-605-098) used in place of anti-fluorescein.
.
**Phagocytosis of bacterial cells with P1Kfl and anti-Fl antibodies.** Vancomycin resistant *E. faecalis* were grown to stationary phase in BHI while shaking (250 rpm) at 37 °C. Bacterial cells from the overnight growth were used to inoculate BHI (1:100) supplemented with 100 µM DADA and incubated at 37 °C with shaking (250 rpm) for 16 h. The bacteria were harvested and washed three times with 1X PBS. The bacteria were resuspended in 50 μL of PBS containing 5 μM of P1Kfl and P1Kfls, 10 % (v/v) FBS, and 0.03 mg/mL IgG fraction monoclonal mouse anti-fluorescein (FITC) (Jackson Immuno Research #200-002-037) and incubated at 37°C for 30 min with shaking (250 rpm) protected from light. The opsonized bacteria were washed with 1× PBS. J774A.1 cells were cultured as described prior. On the day of the experiment, J774A.1 cells were washed with 1X PBS by centrifuging 5 min at 1,000 rpm. The washed J774A cells were then mixed with opsonized *E. faecalis* (MOI 100) in DMEM + 10% FBS containing no antibiotics. The cell mixture was then incubated at 37 °C for 30 min to induce phagocytosis. The cell mixture was centrifuged for 5 min at 1,000 rpm and media was replaced with DMEM + 10% FBS + 300 μg/mL gentamycin and incubated at 4°C for 30 min. The cells were washed three times with 1X PBS and fixed for 30 min with 4% formaldehyde in 1X PBS. Samples were then analyzed by flow cytometry as described above.

### Scheme S1. Synthesis of P1fl

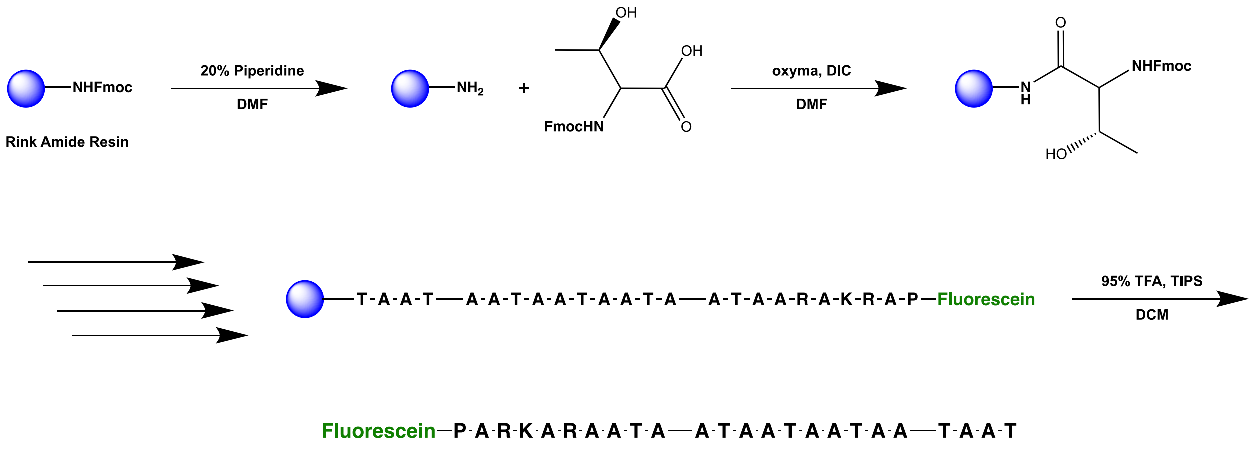


P1fl

A 25 mL vessel of a CEM Discover Bio Manual Peptide Synthesizer was charged with 500 mg (0.27 mmol) of Rink Amide resin. The Fmoc group was removed using a 20% piperidine in DMF solution (10 mL). Using Synergy software, the deprotection protocol was run. The piperidine solution was drained and the resin was washed with DMF (4 x 10 mL). Fmoc-Thr(tBu)-OH (5 eq, 321 mg, 1.35 mmol), 1 M Oxyma (5 eq, 1.35 mmol), and 1 M DIC (5 eq, 1.35 mmol) in DMF (10 mL) were added to the reaction flask and the coupling protocol was run. The amino acid solution was drained, and the resin was washed with DMF (2 x 10 mL). The Fmoc removal and coupling procedure was repeated as before using the same equivalencies for the following amino acids: AATAATAATAATAATAARAKRAP. The Fmoc group of Proline was removed, and resin was transferred to a 25 mL synthetic peptide vessel and coupled 5,6-carboxyfluorescein (3 eq, 304 mg, 0.81 mmol), HBTU (3 eq, 301 mg, 0.81 mmol), and DIEA (6 eq, 281 uL, 1.62 mmol) in DMF (15 mL) shaking over-night. The resin was washed with DCM and MeOH (3 x 15 mL each). To remove the peptide from resin, a TFA cocktail solution (95 % TFA, 2.5% TIPS, and 2.5 % DCM) was added to the resin with agitation for 2 hours protected from light. The resin was filtered and resulting solution was concentrated *in vacuo*. The peptide was trituated with cold diethyl ether and purified using reverse phase HPLC using $H_2O$/MeOH to yield **P1fl**. The sample was analyzed for purity using a Waters 1525 Binary HPLC Pump using a Phenomenex Luna 5u C8(2) 100A (250 x 4.60 mm) column; gradient elution with $H_2O$/$CH_3CN$. Molecular weight was confirmed using a Shimadzu MALDI-TOF Mass Spectrometer (MALDI-8020).

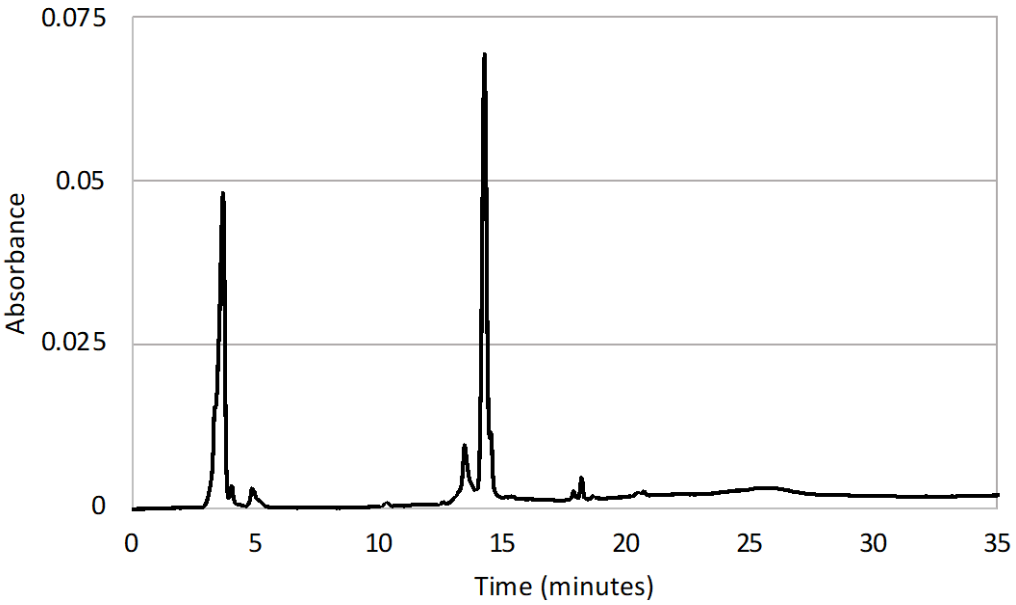


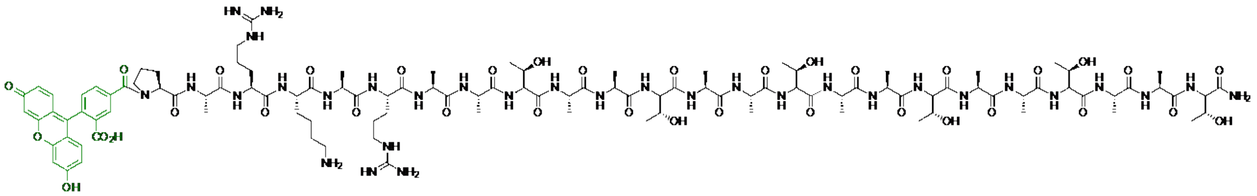

*m/z* calculated: 2514.223, found: 2514.609

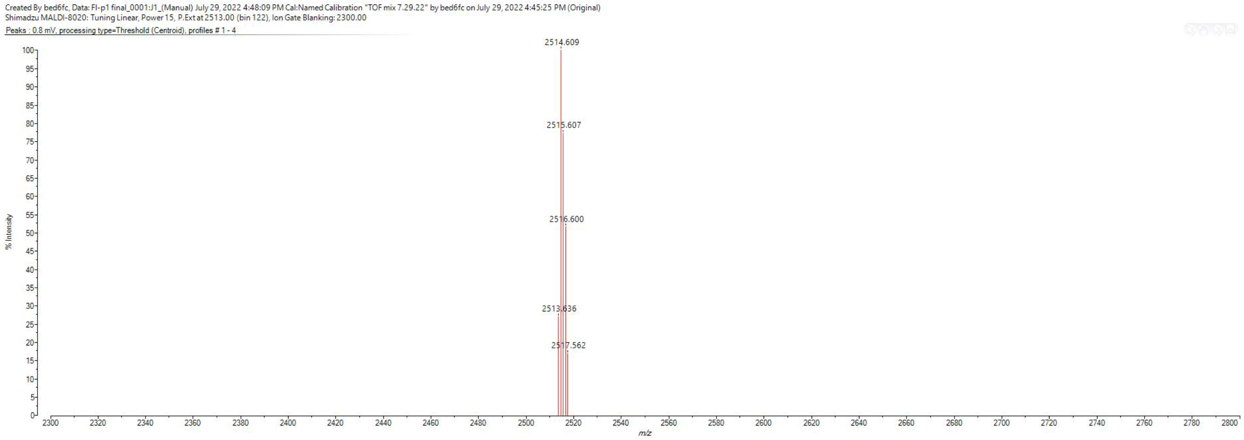

### Scheme S2. Synthesis of P1Kfl

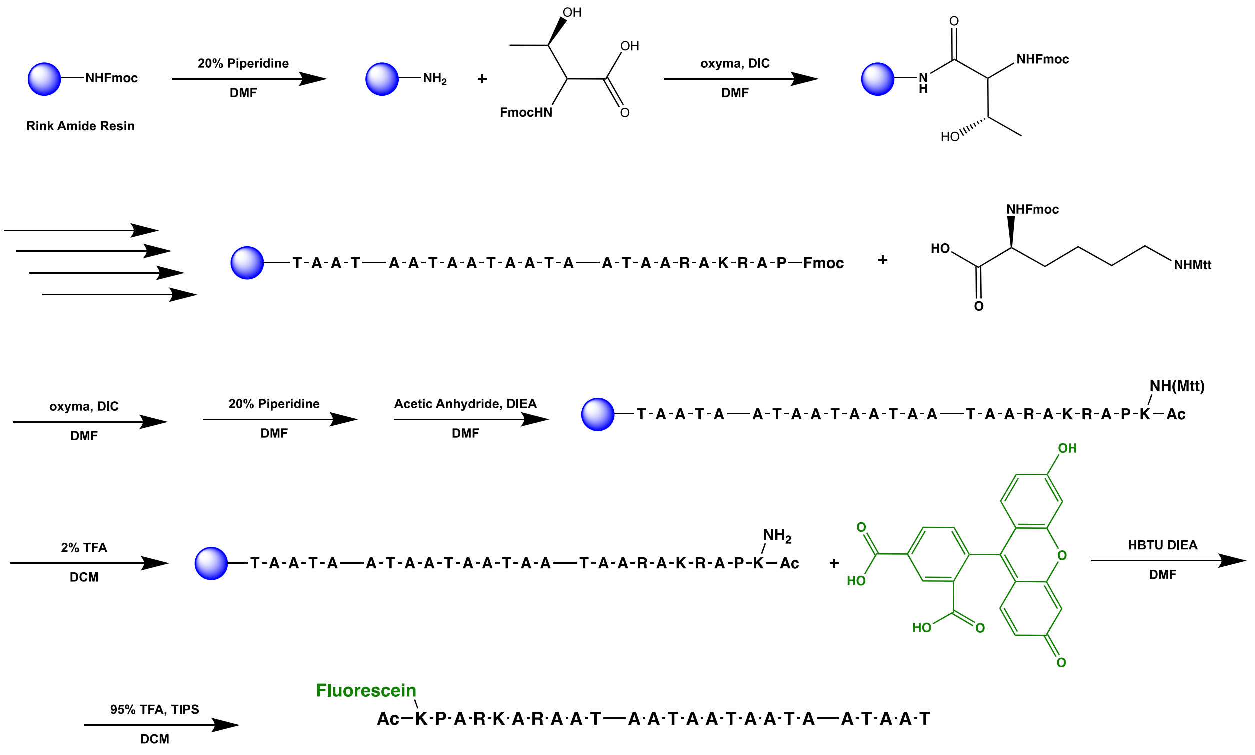


P1Kfl

A 25 mL vessel of a CEM Discover Bio Manual Peptide Synthesizer was charged with 500 mg (0.27 mmol) of Rink Amide resin. The Fmoc group was removed using a 20% piperidine in DMF solution (10 mL). Using Synergy software, the deprotection protocol was run. The piperidine solution was drained and the resin was washed with DMF (4 x 10 mL). Fmoc-Thr(tBu)-OH (5 eq, 321 mg, 1.35 mmol), 1 M Oxyma (5 eq, 1.35 mmol), and 1 M DIC (5 eq, 1.35 mmol) in DMF (10 mL) were added to the reaction flask and the coupling protocol was run. The amino acid solution was drained, and the resin was washed with DMF (2 x 10 mL). The Fmoc removal and coupling procedure was repeated as before using the same equivalencies for the following amino acids: AATAATAATAATAATAARAKRAP. The Fmoc group of Proline was removed and Fmoc-L-Lysine(Mtt)-OH (5 eq, 505 mg, 1.35 mmol) was coupled. The resin was transferred to a 25 mL synthetic vessel and the N-terminus of the peptide was acetylated by removing the Fmoc group and then agitating the resin for 1 hour with a solution of 5% acetic anhydride (0.5 mL), 8.5% DIEA (0.85 mL), and 86.5% DMF (8.65 mL). The Mtt protecting group of L-Lysine(Mtt)-OH was removed by adding 10 mL of a TFA cocktail solution (1% TFA, 2% TIPS in DCM) to the resin and agitating for 10 minutes protected from light. The solution was drained, and this procedure was repeated five additional times. The solution was then drained, rinsed with DMF, and washed as previously described. Upon Mtt removal, 5,6-carboxyfluorescein (3 eq, 304 mg, 0.81 mmol), HBTU (3 eq, 301 mg, 0.81 mmol), and DIEA (6 eq, 281 uL, 1.62 mmol) in DMF (15 mL) was agitated with the resin over-night. The resin was washed with DCM and MeOH (3 x 15 mL each). To remove the peptide from resin, a TFA cocktail solution (95 % TFA, 2.5% TIPS, and 2.5 % DCM) was added to the resin with agitation for 2 hours protected from light. The resin was filtered and resulting solution was concentrated *in vacuo*. The peptide was trituated with cold diethyl ether and purified using reverse phase HPLC using $H_2O$/MeOH to yield **P1Kfl**. The sample was analyzed for purity using a Waters 1525 Binary HPLC Pump using a Phenomenex Luna 5u C8(2) 100A (250 x 4.60 mm) column; gradient elution with $H_2O/CH_3CN$. Molecular weight was confirmed using a Shimadzu MALDI-TOF Mass Spectrometer (MALDI-8020).

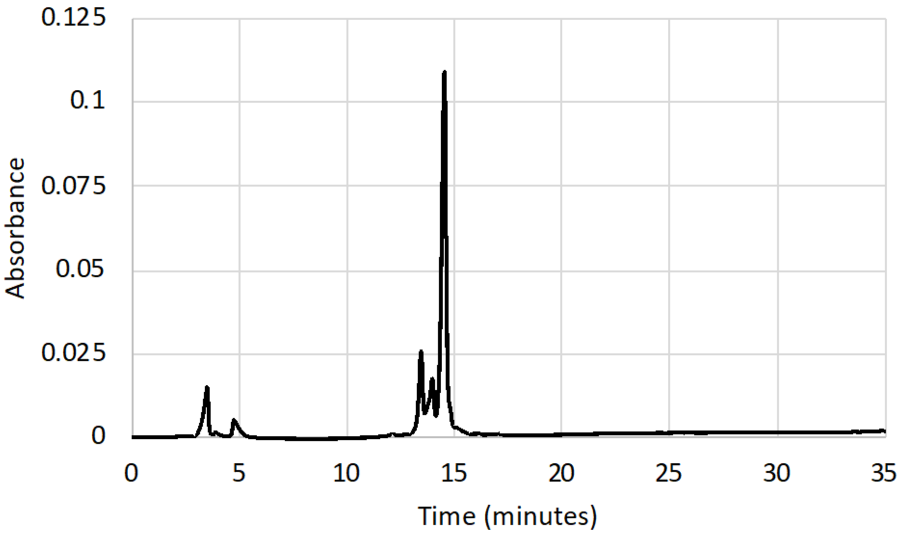


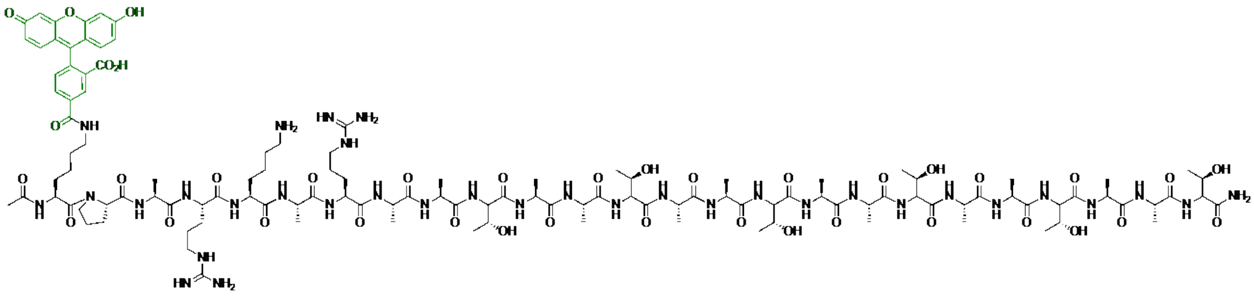

*m/z* calculated: 2684.339, found: 2684.046

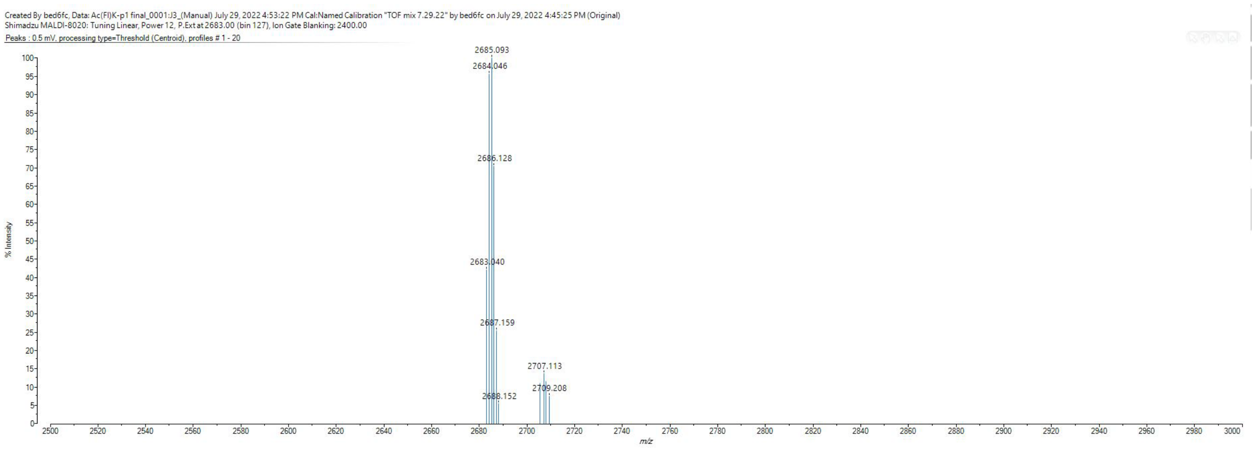

#### Scheme S3. Synthesis of P1fls

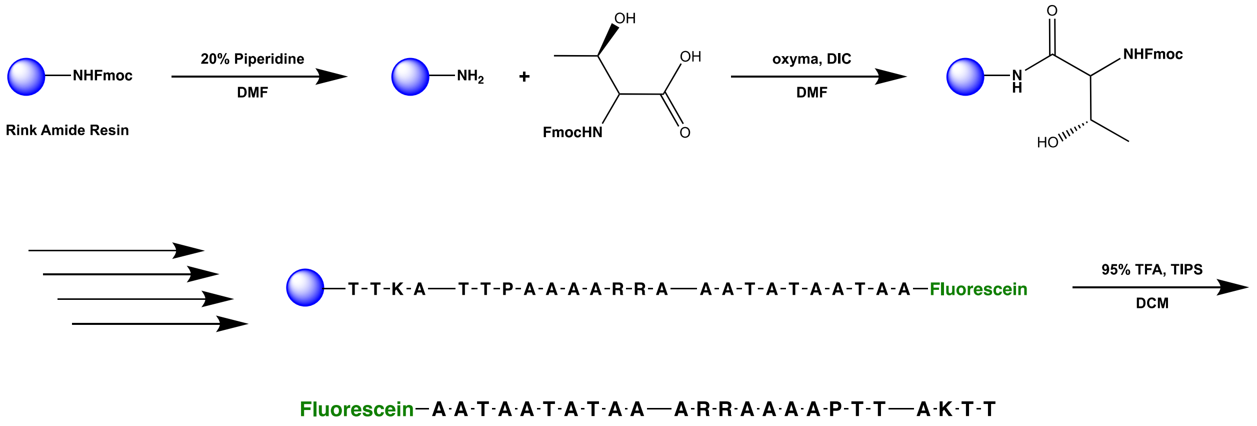


**P1fls**

A 25 mL vessel of a CEM Discover Bio Manual Peptide Synthesizer was charged with 500 mg (0.27 mmol) of Rink Amide resin. The Fmoc group was removed using a 20% piperidine in DMF solution (10 mL). Using Synergy software, the deprotection protocol was run. The piperidine solution was drained and the resin was washed with DMF (4 x 10 mL). Fmoc-Thr(tBu)-OH (5 eq, 321 mg, 1.35 mmol), 1 M Oxyma (5 eq, 1.35 mmol), and 1 M DIC (5 eq, 1.35 mmol) in DMF (10 mL) were added to the reaction flask and the coupling protocol was run. The amino acid solution was drained, and the resin was washed with DMF (2 x 10 mL). The Fmoc removal and coupling procedure was repeated as before using the same equivalencies for the following amino acids: TKATTPAAAARRAAATATAATAA. The Fmoc group of Alanine was removed, and resin was transferred to a 25 mL synthetic peptide vessel and coupled 5,6-carboxyfluorescein (3 eq, 304 mg, 0.81 mmol), HBTU (3 eq, 301 mg, 0.81 mmol), and DIEA (6 eq, 281 uL, 1.62 mmol) in DMF (15 mL) shaking over-night. The resin was washed with DCM and MeOH (3 x 15 mL each). To remove the peptide from resin, a TFA cocktail solution (95 % TFA, 2.5% TIPS, and 2.5 % DCM) was added to the resin with agitation for 2 hours protected from light. The resin was filtered and resulting solution was concentrated *in vacuo*. The peptide was trituated with cold diethyl ether and purified using reverse phase HPLC using $H_2O$/MeOH to yield **P1fls**. The sample was analyzed for purity using a Waters 1525 Binary HPLC Pump using a Phenomenex Luna 5u C8(2) 100A (250 x 4.60 mm) column; gradient elution with $H_2O$/$CH_3CN$. Molecular weight was confirmed using a Shimadzu MALDI-TOF Mass Spectrometer (MALDI-8020).

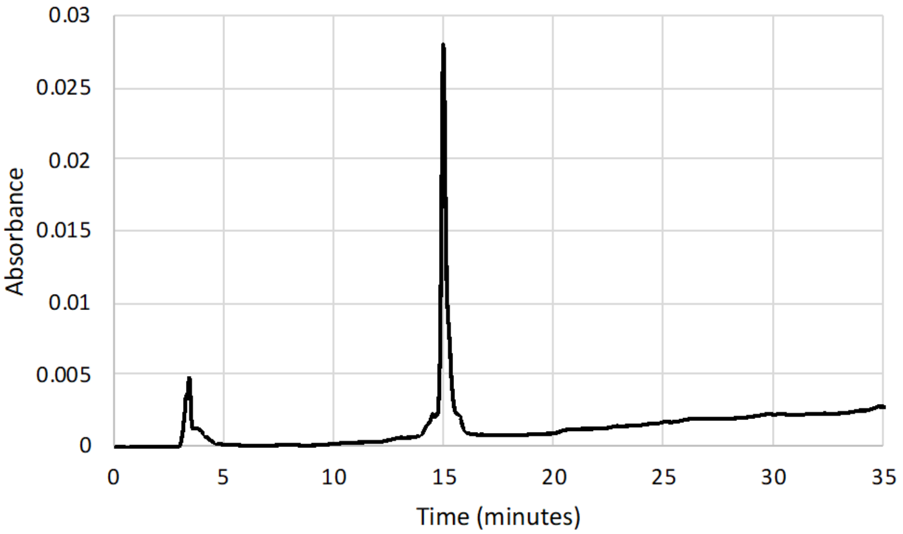


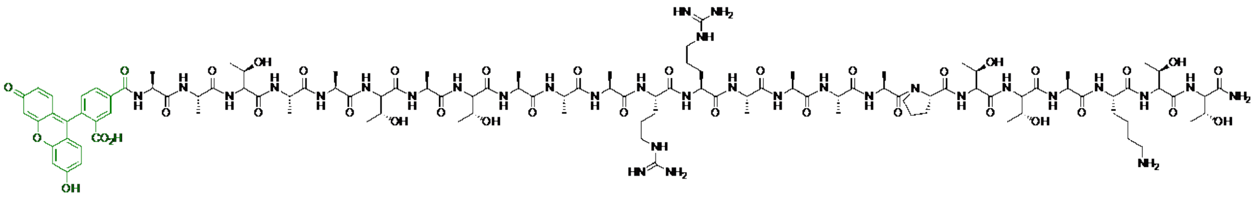

*m/z* calculated: 2544.249, found: 2544.506

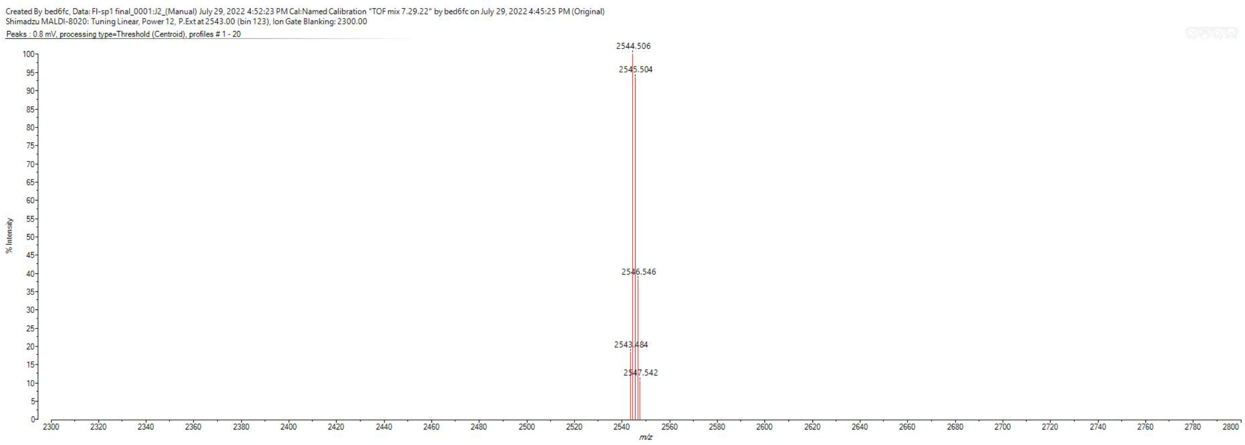

### Scheme S4. Synthesis of P1Kfls

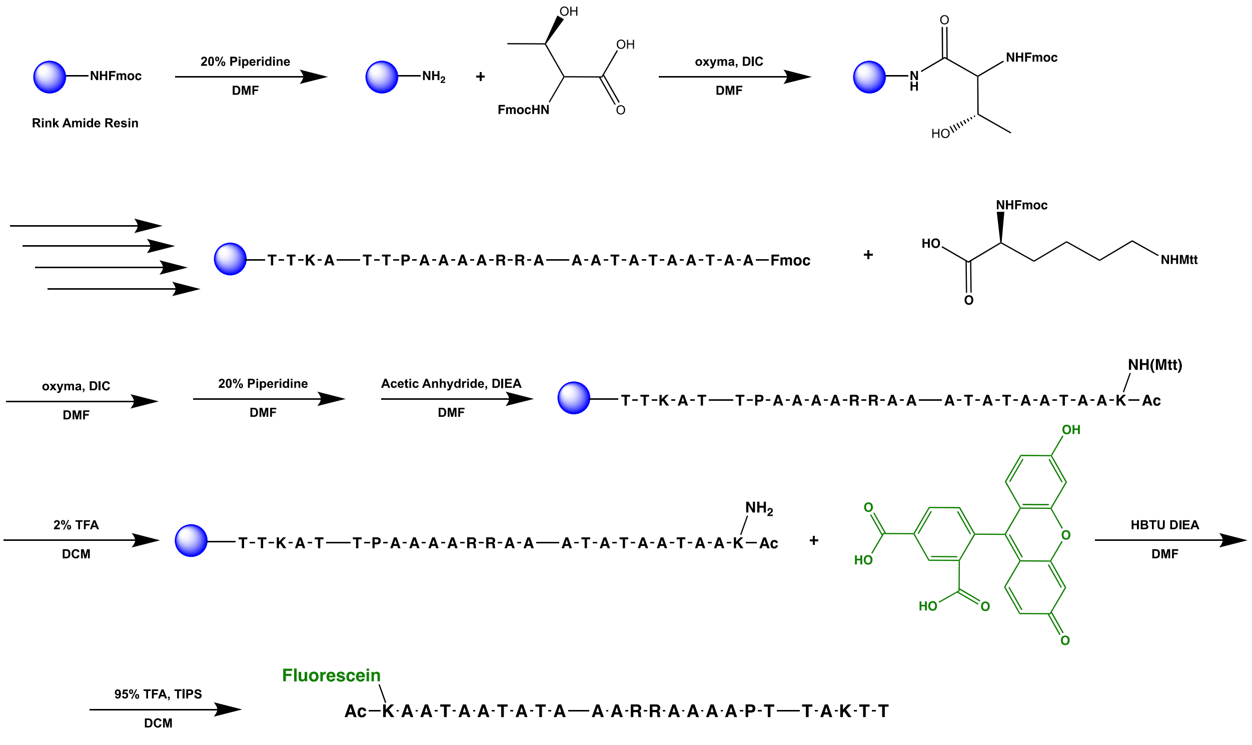


A 25 mL vessel of a CEM Discover Bio Manual Peptide Synthesizer was charged with 500 mg (0.27 mmol) of Rink Amide resin. The Fmoc group was removed using a 20% piperidine in DMF solution (10 mL). Using Synergy software, the deprotection protocol was run. The piperidine solution was drained and the resin was washed with DMF (4 x 10 mL). Fmoc-Thr(tBu)-OH (5 eq, 321 mg, 1.35 mmol), 1 M Oxyma (5 eq, 1.35 mmol), and 1 M DIC (5 eq, 1.35 mmol) in DMF (10 mL) were added to the reaction flask and the coupling protocol was run. The amino acid solution was drained, and the resin was washed with DMF (2 x 10 mL). The Fmoc removal and coupling procedure was repeated as before using the same equivalencies for the following amino acids: TKATTPAAAARRAAATATAATAA. The Fmoc group of Alanine was removed and Fmoc-L-Lysine(Mtt)-OH (5 eq, 505 mg, 1.35 mmol) was coupled. The resin was transferred to a 25 mL synthetic vessel and the N-terminus of the peptide was acetylated by removing the Fmoc group and then agitating the resin for 1 hour with a solution of 5% acetic anhydride (0.5 mL), 8.5% DIEA (0.85 mL), and 86.5% DMF (8.65 mL). The Mtt protecting group of L-Lysine(Mtt)-OH was removed by adding 10 mL of a TFA cocktail solution (1% TFA, 2% TIPS in DCM) to the resin and agitating for 10 minutes protected from light. The solution was drained, and this procedure was repeated five additional times. The solution was then drained, rinsed with DMF, and washed as previously described. Upon Mtt removal, 5,6-carboxyfluorescein (3 eq, 304 mg, 0.81 mmol), HBTU (3 eq, 301 mg, 0.81 mmol), and DIEA (6 eq, 281 uL, 1.62 mmol) in DMF (15 mL) was agitated with the resin over-night. The resin was washed with DCM and MeOH (3 x 15 mL each). To remove the peptide from resin, a TFA cocktail solution (95 % TFA, 2.5% TIPS, and 2.5 % DCM) was added to the resin with agitation for 2 hours protected from light. The resin was filtered and resulting solution was concentrated *in vacuo*. The peptide was triturated with cold diethyl ether and purified using reverse phase HPLC using $H_2O$/MeOH to yield **P1Kfls**. The sample was analyzed for purity using a Waters 1525 Binary HPLC Pump using a Phenomenex Luna 5u C8(2) 100A (250 x 4.60 mm) column; gradient elution with $H_2O$/$CH_3CN$. Molecular weight was confirmed using a Shimadzu MALDI-TOF Mass Spectrometer (MALDI-8020).

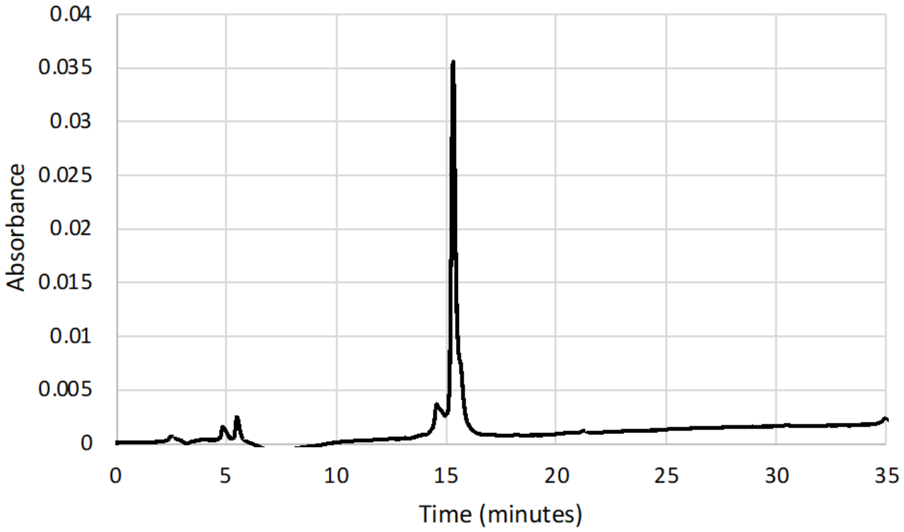


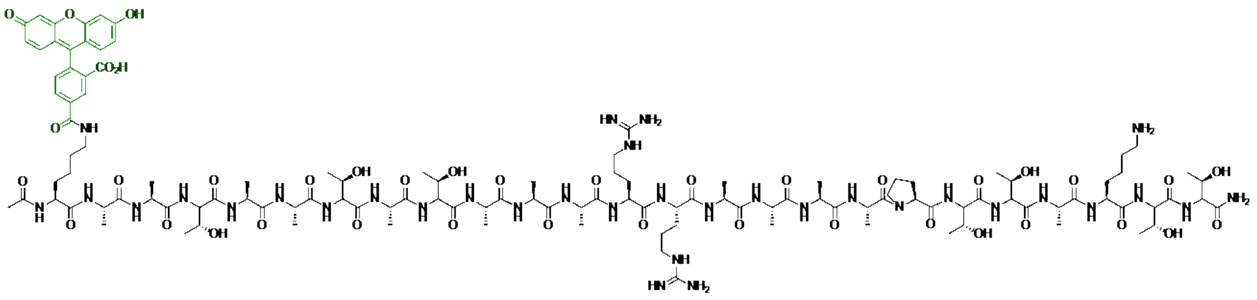

*m/z* calculated: 2714.350, found: 2714.533

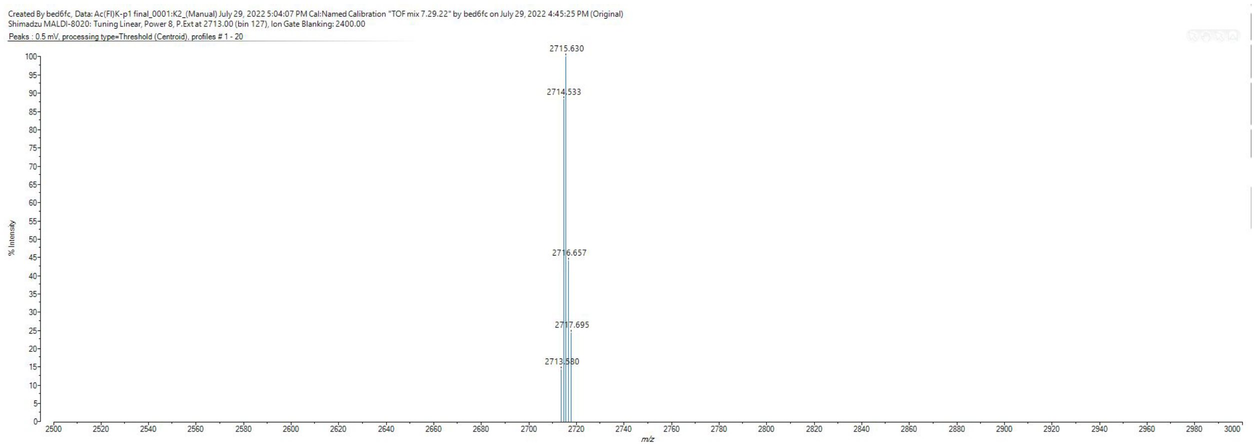

### Scheme S5. Synthesis of P1dnp

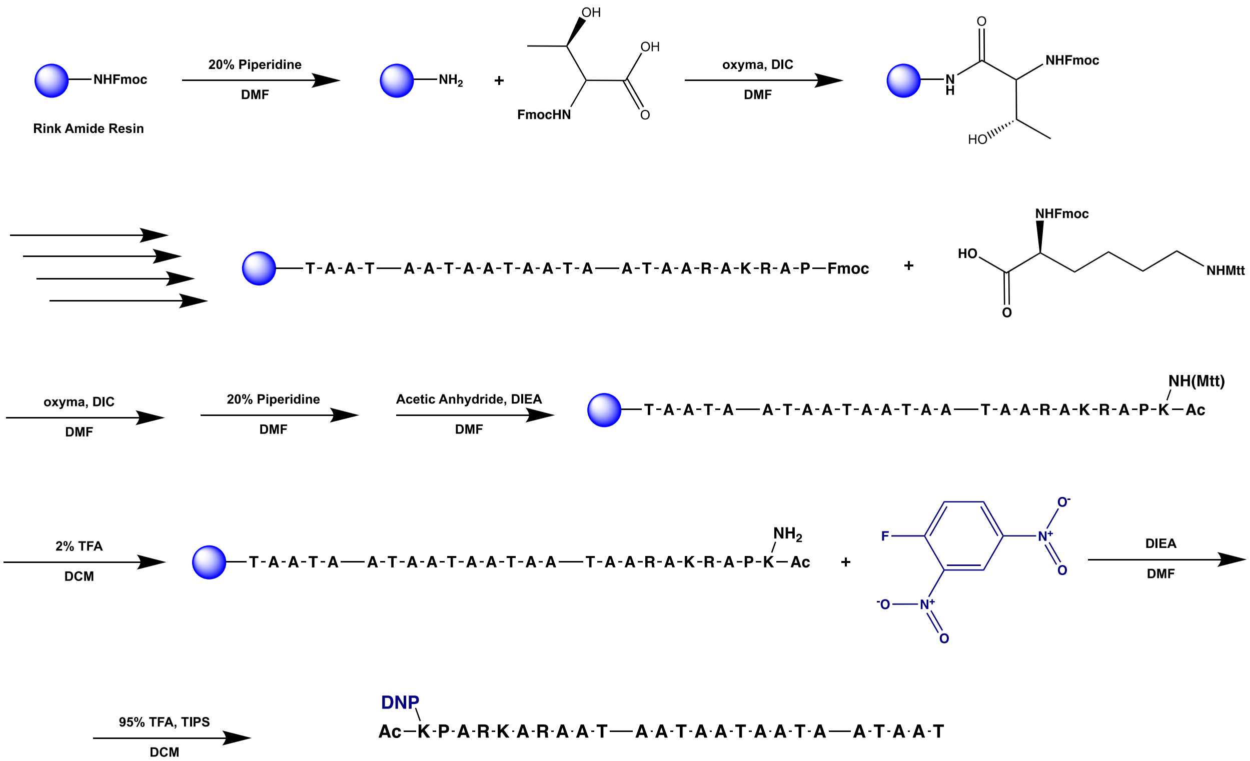


**P1dnp**

A 25 mL vessel of a CEM Discover Bio Manual Peptide Synthesizer was charged with 500 mg (0.27 mmol) of Rink Amide resin. The Fmoc group was removed using a 20% piperidine in DMF solution (10 mL). Using Synergy software, the deprotection protocol was run. The piperidine solution was drained and the resin was washed with DMF (4 x 10 mL). Fmoc-Thr(tBu)-OH (5 eq, 321 mg, 1.35 mmol), 1 M Oxyma (5 eq, 1.35 mmol), and 1 M DIC (5 eq, 1.35 mmol) in DMF (10 mL) were added to the reaction flask and the coupling protocol was run. The amino acid solution was drained, and the resin was washed with DMF (2 x 10 mL). The Fmoc removal and coupling procedure was repeated as before using the same equivalencies for the following amino acids: AATAATAATAATAATAARAKRAP. The Fmoc group of Proline was removed and Fmoc-L-Lysine(Mtt)-OH (5 eq, 505 mg, 1.35 mmol) was coupled. The resin was transferred to a 25 mL synthetic vessel and the N-terminus of the peptide was acetylated by removing the Fmoc group and then agitating the resin for 1 hour with a solution of 5% acetic anhydride (0.5 mL), 8.5% DIEA (0.85 mL), and 86.5% DMF (8.65 mL). The Mtt protecting group of L-Lysine(Mtt)-OH was removed by adding 10 mL of a TFA cocktail solution (1% TFA, 2% TIPS in DCM) to the resin and agitating for 10 minutes protected from light. The solution was drained, and this procedure was repeated five additional times. The solution was then drained, rinsed with DMF, and washed as previously described. Upon Mtt removal, 2,4-dinitrofluorobenzene (3 eq, 100 uL, 1.35 mmol) and DIEA (6 eq, 467 uL, 1.62 mmol) in DMF (15 mL) was agitated with the resin for 2 hours protected from light. The resin was washed with DCM and MeOH (3 x 15 mL each). To remove the peptide from resin, a TFA cocktail solution (95 % TFA, 2.5% TIPS, and 2.5 % DCM) was added to the resin with agitation for 2 hours protected from light. The resin was filtered and resulting solution was concentrated *in vacuo*. The peptide was trituated with cold diethyl ether and purified using reverse phase HPLC using $H_2O$/MeOH to yield **P1dnp**. The sample was analyzed for purity using a Waters 1525 Binary HPLC Pump using a Phenomenex Luna 5u C8(2) 100A (250 x 4.60 mm) column; gradient elution with $H_2O$/$CH_3CN$. Molecular weight was confirmed using a Shimadzu MALDI-TOF Mass Spectrometer (MALDI-8020).

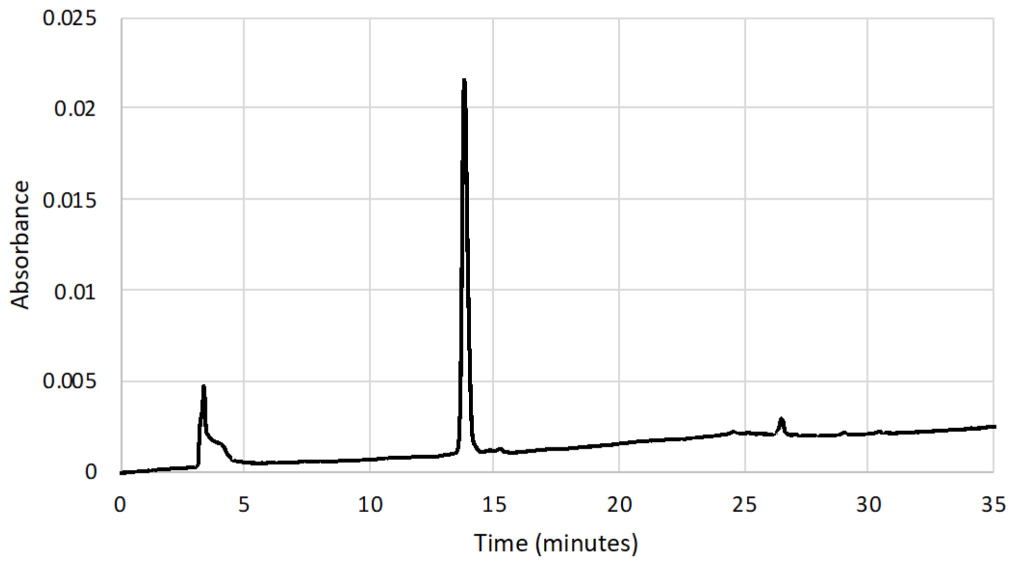


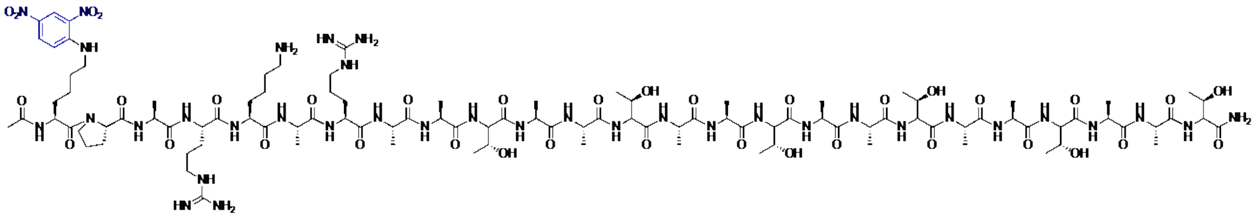

*m/z* calculated: 2492.293, found: 2492.414

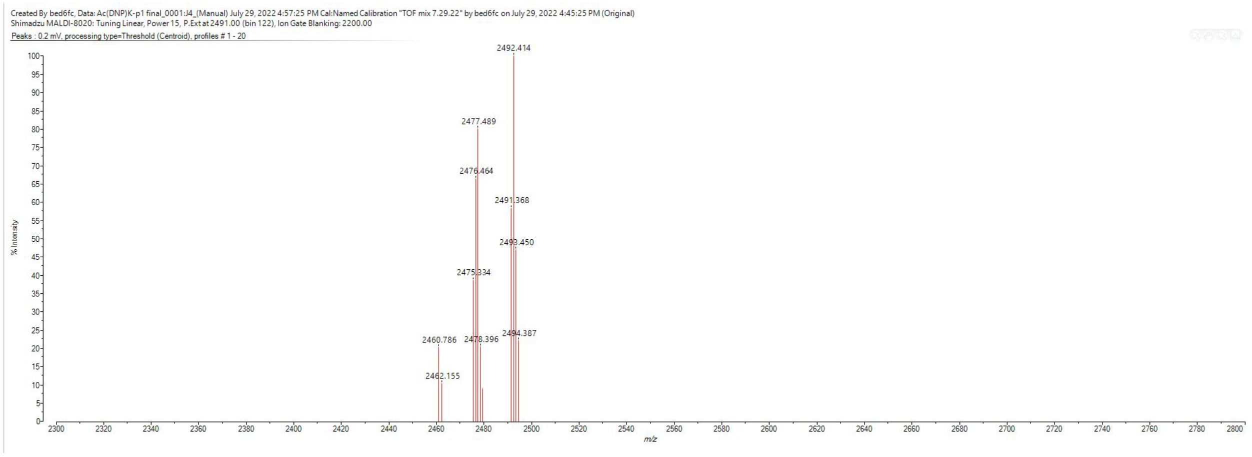

### Scheme S6. Synthesis of P1dnps

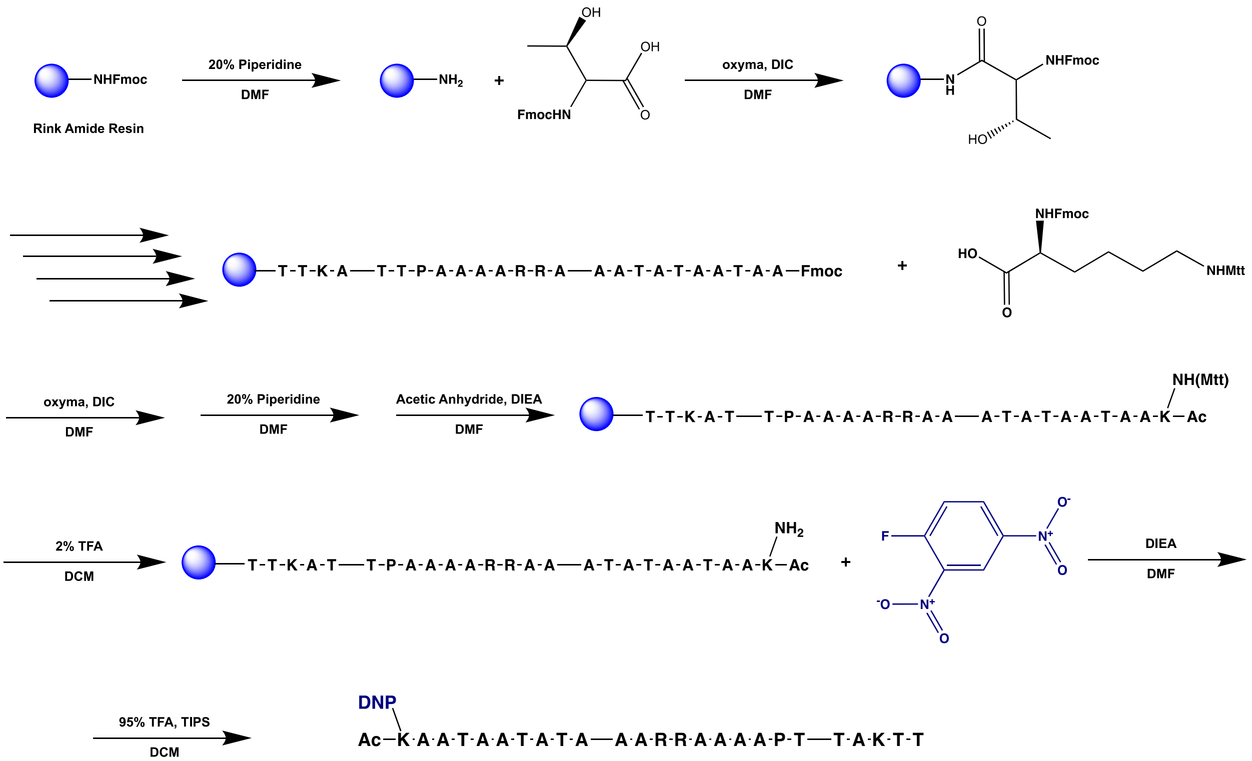


**P1dnps**

A 25 mL vessel of a CEM Discover Bio Manual Peptide Synthesizer was charged with 500 mg (0.27 mmol) of Rink Amide resin. The Fmoc group was removed using a 20% piperidine in DMF solution (10 mL). Using Synergy software, the deprotection protocol was run. The piperidine solution was drained and the resin was washed with DMF (4 x 10 mL). Fmoc-Thr(tBu)-OH (5 eq, 321 mg, 1.35 mmol), 1 M Oxyma (5 eq, 1.35 mmol), and 1 M DIC (5 eq, 1.35 mmol) in DMF (10 mL) were added to the reaction flask and the coupling protocol was run. The amino acid solution was drained, and the resin was washed with DMF (2 x 10 mL). The Fmoc removal and coupling procedure was repeated as before using the same equivalencies for the following amino acids: TKATTPAAAARRAAATATAATAA. The Fmoc group of Alanine was removed and Fmoc-L-Lysine(Mtt)-OH (5 eq, 505 mg, 1.35 mmol) was coupled. The resin was transferred to a 25 mL synthetic vessel and the N-terminus of the peptide was acetylated by removing the Fmoc group and then agitating the resin for 1 hour with a solution of 5% acetic anhydride (0.5 mL), 8.5% DIEA (0.85 mL), and 86.5% DMF (8.65 mL). The Mtt protecting group of L-Lysine(Mtt)-OH was removed by adding 10 mL of a TFA cocktail solution (1% TFA, 2% TIPS in DCM) to the resin and agitating for 10 minutes protected from light. The solution was drained, and this procedure was repeated five additional times. The solution was then drained, rinsed with DMF, and washed as previously described. Upon Mtt removal, 2,4-dinitrofluorobenzene (3 eq, 100 uL, 1.35 mmol) and DIEA (6 eq, 467 uL, 1.62 mmol) in DMF (15 mL) was agitated with the resin for 2 hours protected from light. The resin was washed with DCM and MeOH (3 x 15 mL each). To remove the peptide from resin, a TFA cocktail solution (95 % TFA, 2.5% TIPS, and 2.5 % DCM) was added to the resin with agitation for 2 hours protected from light. The resin was filtered and resulting solution was concentrated *in vacuo*. The peptide was trituated with cold diethyl ether and purified using reverse phase HPLC using $H_2O$/MeOH to yield **P1dnps**. The sample was analyzed for purity using a Waters 1525 Binary HPLC Pump using a Phenomenex Luna 5u C8(2) 100A (250 x 4.60 mm) column; gradient elution with $H_2O/CH_3CN$. Molecular weight was confirmed using a Shimadzu MALDI-TOF Mass Spectrometer (MALDI-8020).

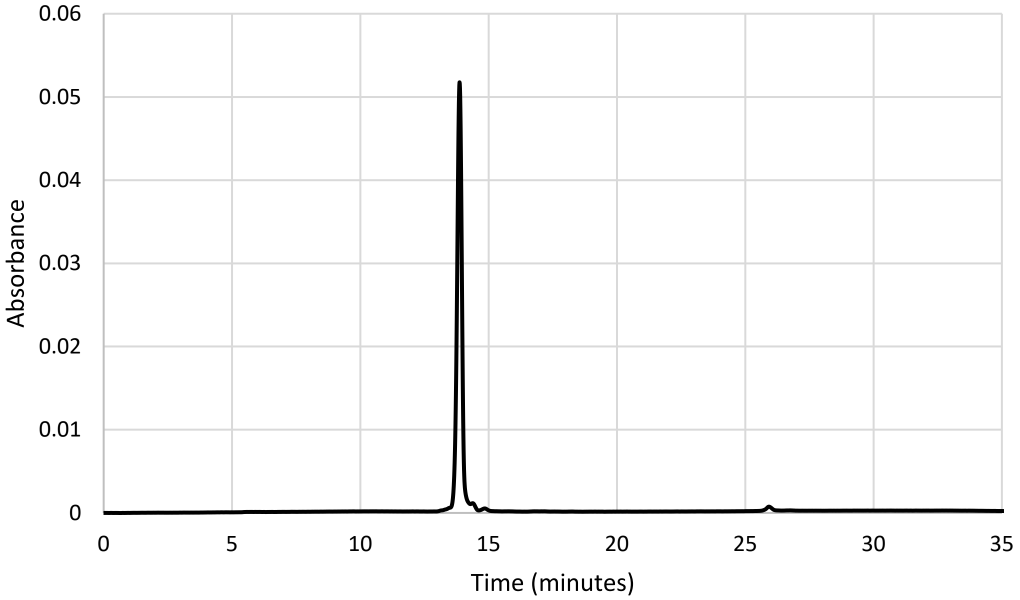


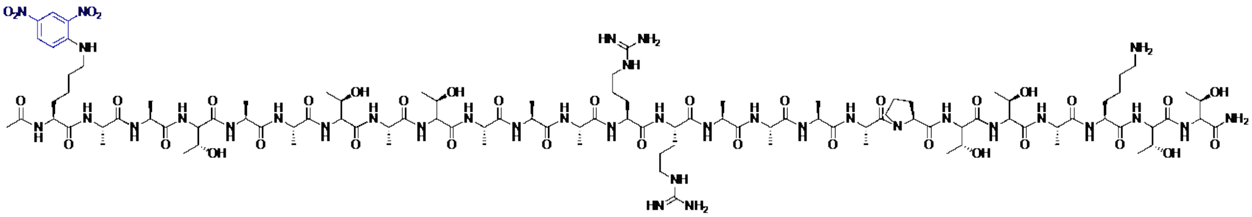

*m/z* calculated: 2522.303, found: 2522.186

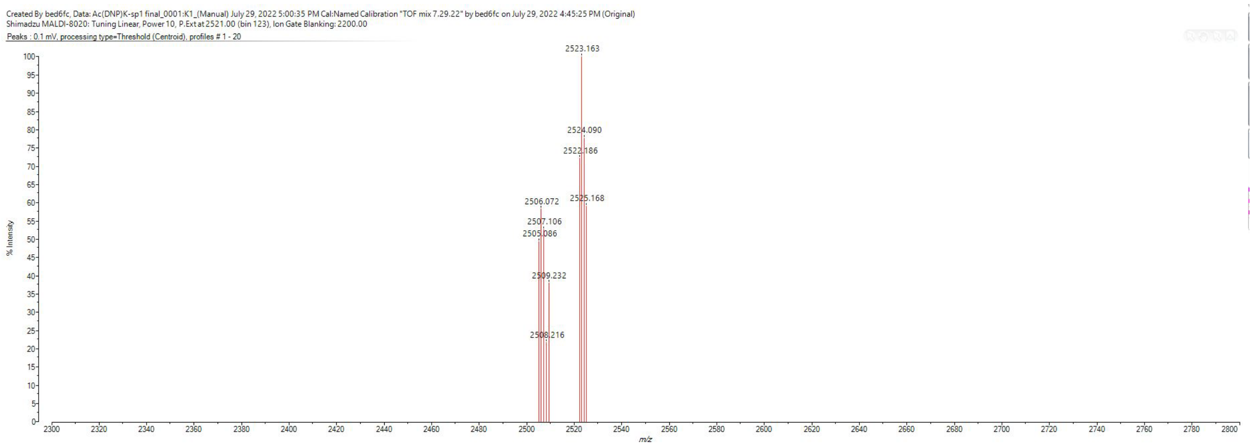

### Scheme S7. Synthesis of P1tam

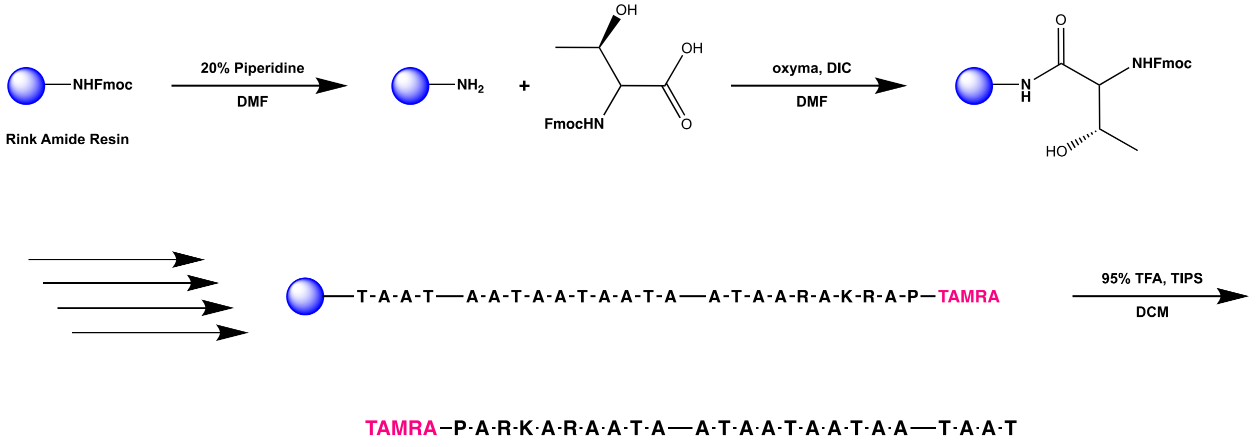


P1tam

A 25 mL vessel of a CEM Discover Bio Manual Peptide Synthesizer was charged with 500 mg (0.27 mmol) of Rink Amide resin. The Fmoc group was removed using a 20% piperidine in DMF solution (10 mL). Using Synergy software, the deprotection protocol was run. The piperidine solution was drained and the resin was washed with DMF (4 x 10 mL). Fmoc-Thr(tBu)-OH (5 eq, 321 mg, 1.35 mmol), 1 M Oxyma (5 eq, 1.35 mmol), and 1 M DIC (5 eq, 1.35 mmol) in DMF (10 mL) were added to the reaction flask and the coupling protocol was run. The amino acid solution was drained, and the resin was washed with DMF (2 x 10 mL). The Fmoc removal and coupling procedure was repeated as before using the same equivalencies for the following amino acids: AATAATAATAATAATAARAKRAP. The Fmoc group of Proline was removed, and resin was transferred to a 25 mL synthetic peptide vessel and coupled 5,6-carboxy-tetramethylrhodamine (3 eq, 304 mg, 0.81 mmol), HBTU (3 eq, 301 mg, 0.81 mmol), and DIEA (6 eq, 281 uL, 1.62 mmol) in DMF (15 mL) shaking over-night. The resin was washed with DCM and MeOH (3 x 15 mL each). To remove the peptide from resin, a TFA cocktail solution (95 % TFA, 2.5% TIPS, and 2.5 % DCM) was added to the resin with agitation for 2 hours protected from light. The resin was filtered and resulting solution was concentrated *in vacuo*. The peptide was trituated with cold diethyl ether and purified using reverse phase HPLC using $H_2O$/MeOH to yield **P1tam**. The sample was analyzed for purity using a Waters 1525 Binary HPLC Pump using a Phenomenex Luna 5u C8(2) 100A (250 x 4.60 mm) column; gradient elution with $H_2O/CH_3CN$. Molecular weight was confirmed using a Shimadzu MALDI-TOF Mass Spectrometer (MALDI-8020).

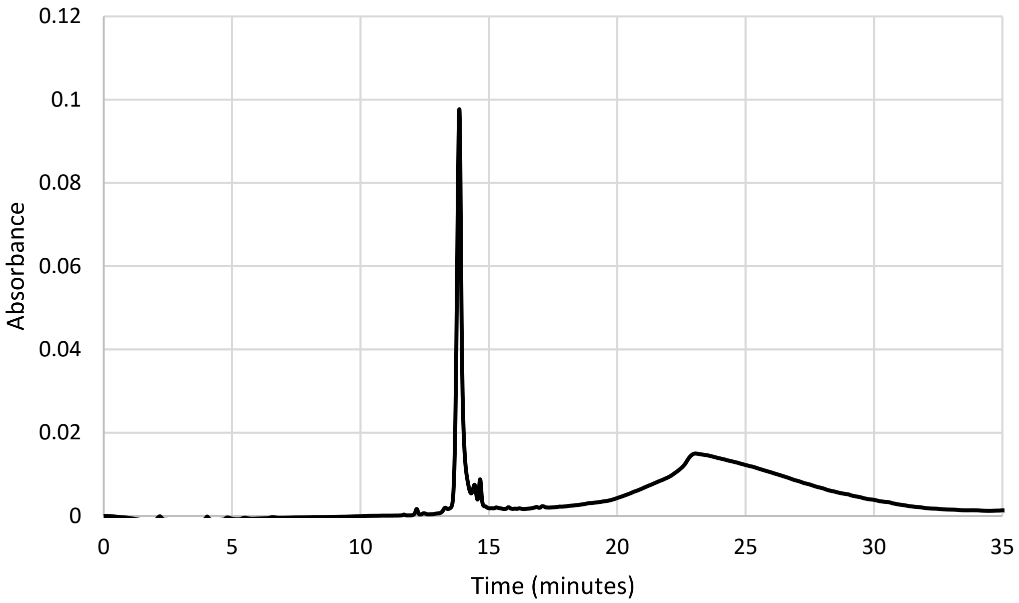


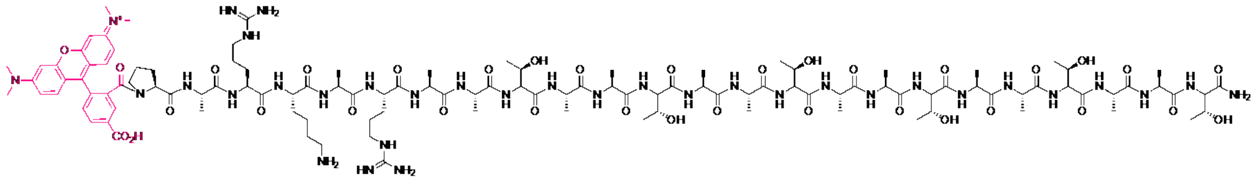

*m/z* calculated: 2569.335, found: 2569.668

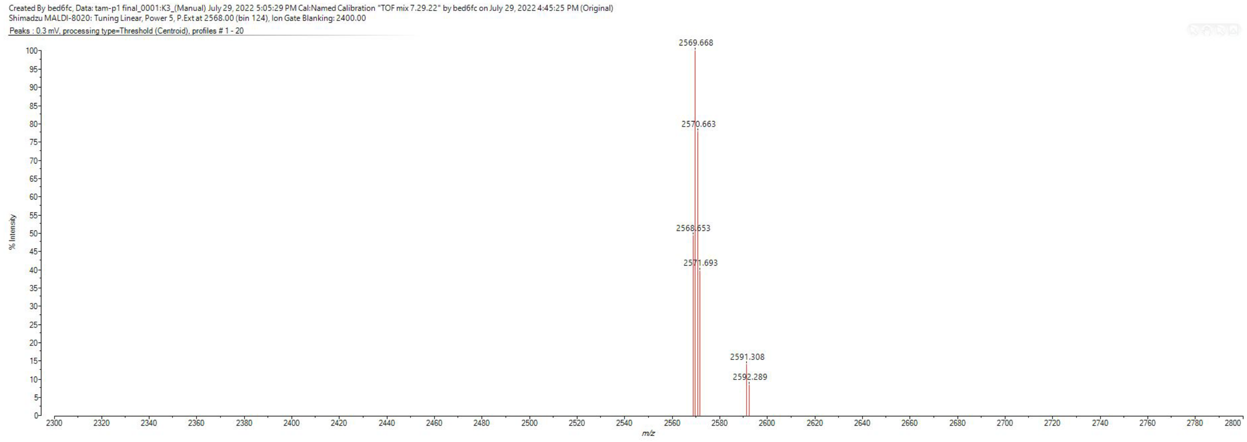
